# CCL19⁺ fibroblasts define a proliferative niche in chronic lymphocytic leukemia

**DOI:** 10.1101/2024.11.15.623868

**Authors:** Antonio P. Ferreira, Shumei Wang, Larysa Poluben, Dominique Morais, Joshua Brandstadter, Eric Perkey, Steven Sotirakos, Mai Drew, Li Pan, Madeleine E. Lemieux, Stacey M Fernandes, David M. Dorfman, Brent Shoji, Sean R. Stowell, Ivan Maillard, Jennifer Brown, Stephen C. Blacklow, Jon C. Aster

## Abstract

Adaptive immune responses occur lymph nodes (LNs) in a microenvironment established by resident stromal cells. LNs are also a site of proliferation of chronic lymphocytic leukemia (CLL), a B cell cancer that alters LN structure in a stereotypic manner. To deeply characterize reactive and CLL LNs, we developed a single-cell RNA sequencing pipeline. We find that proliferation of CLL cells in proliferation centers (PCs), a CLL-specific niche, begins with transient upregulation of *MYC,* subsequent downregulation of which may limit CLL growth. PCs contain a distinct fibroblast population expressing *CCL19* while CLL cells express the CCL19 receptor CCR7, providing a recruitment mechanism for CLL cells to PCs. Using informatic, spatial, and in situ analyses to identify ligand-receptor pairs involving PC CLL cells and nearby immune and stromal cells, we observe that PCs are enriched for macrophages expressing BAFF, the integrin αXβ2 heterodimer, and Galectin9, factors implicated in cell growth, adhesion, and immunosuppression. The most common predicted interactions in PCs involve CD74 and ligands such as MIF, and we find that CD74 blockade consistently inhibits CLL cell growth in culture. Our work highlights key features of the CLL proliferative niche and provides a roadmap for identifying vulnerabilities and new therapeutic strategies.

## Introduction

Lymph nodes (LNs) are specialized organs where B and T lymphocytes interact with antigen-presenting cells to initiate adaptive immune responses^1,2^. These interactions occur within niches organized by stromal fibroblasts, which secrete factors (e.g., chemokines) that recruit immune cells to specific LN compartments to coordinate interactions between lymphocytes and antigen presenting cells^3–5^. Studies have identified several fibroblast subtypes that define different LN areas, such as fibroblastic reticular cells (FRCs)^6^ in T cell zones, marginal reticular cells (MRCs) at boundaries between the capsules and B cell follicles, B cell zone reticular cells proximal to B cell follicles, and follicular dendritic cells (FDCs), which are present in primary and secondary B cell follicles.^4,7,8^

LNs are also the main site of homing and proliferation of chronic lymphocytic leukemia (CLL)^9,10^, a tumor of mature B cells that is the most common leukemia in older adults^11,12^. CLL cells proliferate in loose clusters called proliferation centers (PCs), within which CLL cells enlarge in size and accumulate MYC protein^13^. The focal nature of PCs suggests that local stromal cues orchestrate their formation^14^. Notably, growing CLL cells *ex vivo* has proven challenging, highlighting their likely dependence on factors produced in the nodal microenvironment for growth and survival^15^.

Despite advances in therapy, most CLL cases relapse as drug resistance emerges through multiple mechanisms, including acquired mutations in BTK or PLCγ2 that reactivate B-cell receptor signaling and drive resistance to BTK inhibitors^16^ and mitochondrial reprogramming, which alters apoptotic dependencies and reduces the efficacy of BCL2 inhibitors such as venetoclax^17,18^. Moreover, in a subset of cases persistent CLL transforms into an aggressive B-cell malignancy (Richter transformation), an event that is frequently fatal^19^. The LN microenvironment provides stroma-derived survival signals that protect CLL cells from therapeutic pressure, thereby fostering persistence^9,11^, suggesting that therapies that target components of the proliferative CLL LN microenvironment could complement tumor-directed therapies^20^. To begin to address this need, we deeply characterized CLL cells and their proliferative LN microenvironment using single-cell RNA sequencing (scRNA-seq), spatial transcriptomics, immunofluorescence microscopy, and *ex vivo* spheroid growth assays.

## Results

### Establishment of a pipeline to study human lymph nodes

Because the diagnosis of nodal CLL is typically not established until several days after biopsy, we developed and validated a pipeline (Figure 1A) using viably frozen tissue that produces scRNA-seq data of quality equivalent to that obtained from freshly processed tissue^21^. For initial studies, we selected three LNs involved by reactive hyperplasia or histologically typical CLL (described in Supplemental Table 1), isolated CD45^+^ and CD45^-^ cells as shown in Supplemental Figure 1A, and performed scRNA-seq.

**Figure 1.**
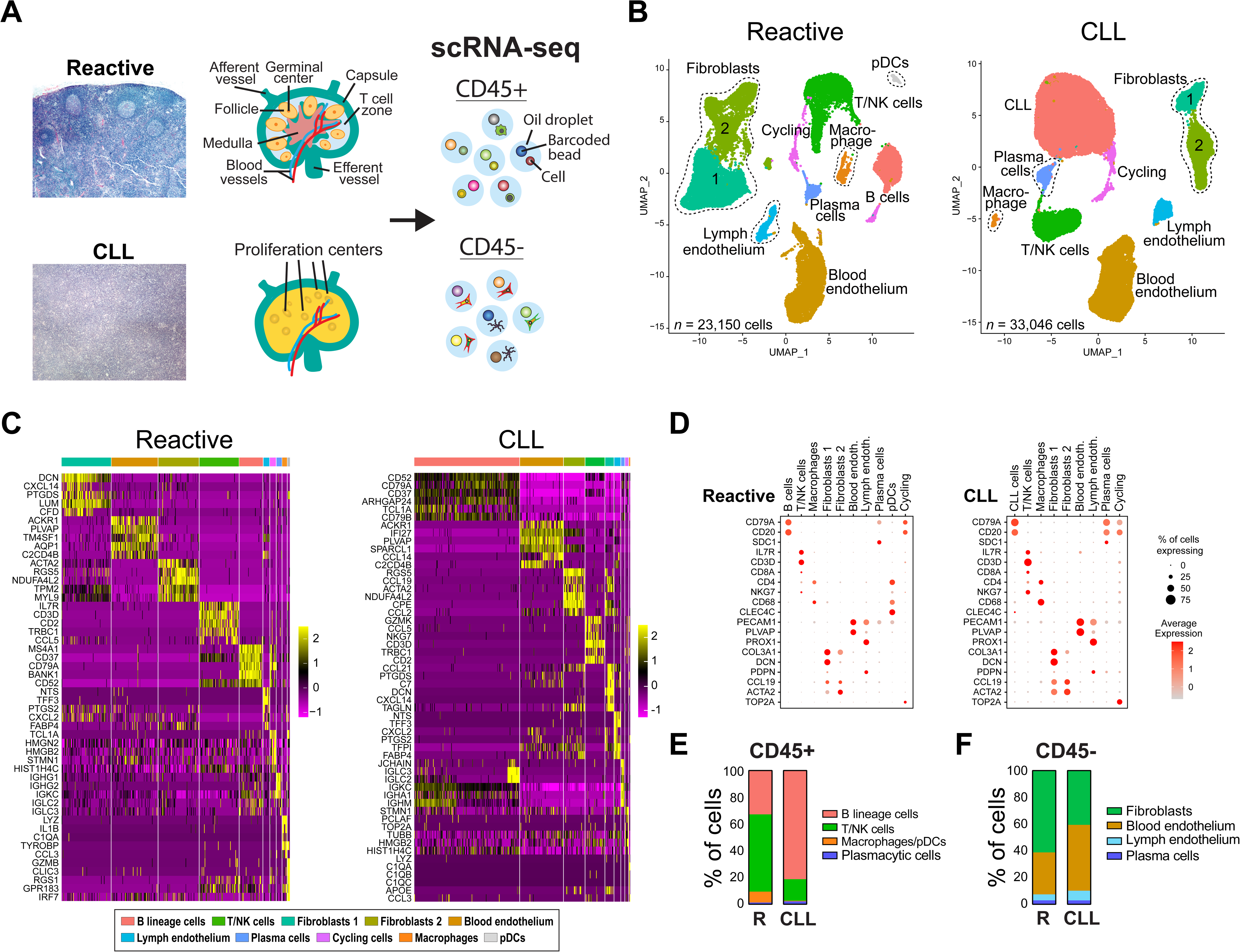
Cell types defined by scRNA-seq in LNs involved by reactive processes and CLL. **A)** scRNA-seq pipeline. Cell suspensions were sorted into CD45^+^ and CD45^-^ fractions and subjected to scRNA-seq. **B)** UMAP plots corresponding to integrated scRNA-seq datasets (3 reactive and 3 CLL LNs). **C)** Heatmaps showing top differentially expressed genes. **D)** Dot plots displaying cluster-specific genes by expression level (shown by color) and percentage of cells (shown by size). **E, F)** Percentages of main CD45^+^ and CD45^-^ cell types identified in reactive (R) and CLL LNs.

We first did a global analysis of all cells, using lineage-specific transcripts (Supplemental Figure 1B and Supplemental Table 2) to identify main cell types (Supplemental Figure 1C-E). In total, high-quality data were obtained from 56,196 cells of diverse types (Supplemental Figure 1E). Noting that some proliferating CLL cells had a level of mitochondrial gene transcripts that would exclude these cells by standard filtering methods, we developed and applied an alternative filter termed N_Mito (see Supplemental Methods) that relies on the ratio of transcripts encoded by mitochondrial and nuclear genes^22^ (see Supplemental Table 2 for nuclear mitochondrial gene list). This alternative filter increased inclusion of proliferating *MYC*^+^ and *MKI67*^+^ cells in downstream analyses (Supplemental Figure 2A-D).

To compare the cellular constituents of CLL and reactive LNs globally, we ran Seurat^23^ followed by Harmony^24^ to integrate scRNA-seq data. This revealed multiple UMAP clusters corresponding to immune and stromal cell types (Figure 1B), as judged by heatmaps (Figure 1C) of differentially expressed genes (Supplemental Table 3) and dot plots displaying expression of lineage-specific markers (Figure 1D). CLL cells were readily distinguished from normal B cells by expression of diagnostically useful markers such as *CD5*, *FCER2* (CD23), and *LEF1* (Supplemental Tables 2-3). As expected, more B lineage cells were recovered from CLL LNs than reactive LNs (Figure 1E) along with more endothelial cells (Figure 1F), consistent with CLL LNs having increased vascularity^25^.

### Characterization of CD45^+^ immune cells in reactive and CLL LNs

Interactions between CLL cells and nodal CD45^+^ immune cells have important pathobiological effects that support CLL cell growth and survival^15^. To study normal and neoplastic immune cells in depth, we first performed a focused integrated analysis of CD45^+^ cells from each sample type at a resolution that identified all expected major cell types (Figure 2A). We found that CLL cells from each sample fell into three subclusters designated CLL1, CLL2, and CLL3 (Figure 2A). Top differentially expressed genes (Figure 2B, Supplemental Table4) in the subclusters included *TXNIP* (CLL subcluster-1), which encodes a protein involved in redox homeostasis^26^; *CD83* (CLL2 subcluster-2), an immunomodulatory factor that is hypothesized to have immunosuppressive effects in CLL patients^27^; and *NME1* and *NME2* (CLL3 subcluster-3), which encode nucleoside diphosphate kinases linked to increased *MYC* expression and altered oxidative phosphorylation in models of Richter transformation^28^. Notably, the CLL3 subcluster also displayed high *MYC* expression, consistent with this cluster being enriched for cells in PCs. Uniquely, CLL case 1 also contained neoplastic B cells with evidence of plasmacytic differentiation, a feature seen in a subset of CLLs^29^. InferCNV analysis confirmed that these plasmacytic cells are clonally related to the accompanying CLL cells (Supplemental Figure 3).

**Figure 2.**
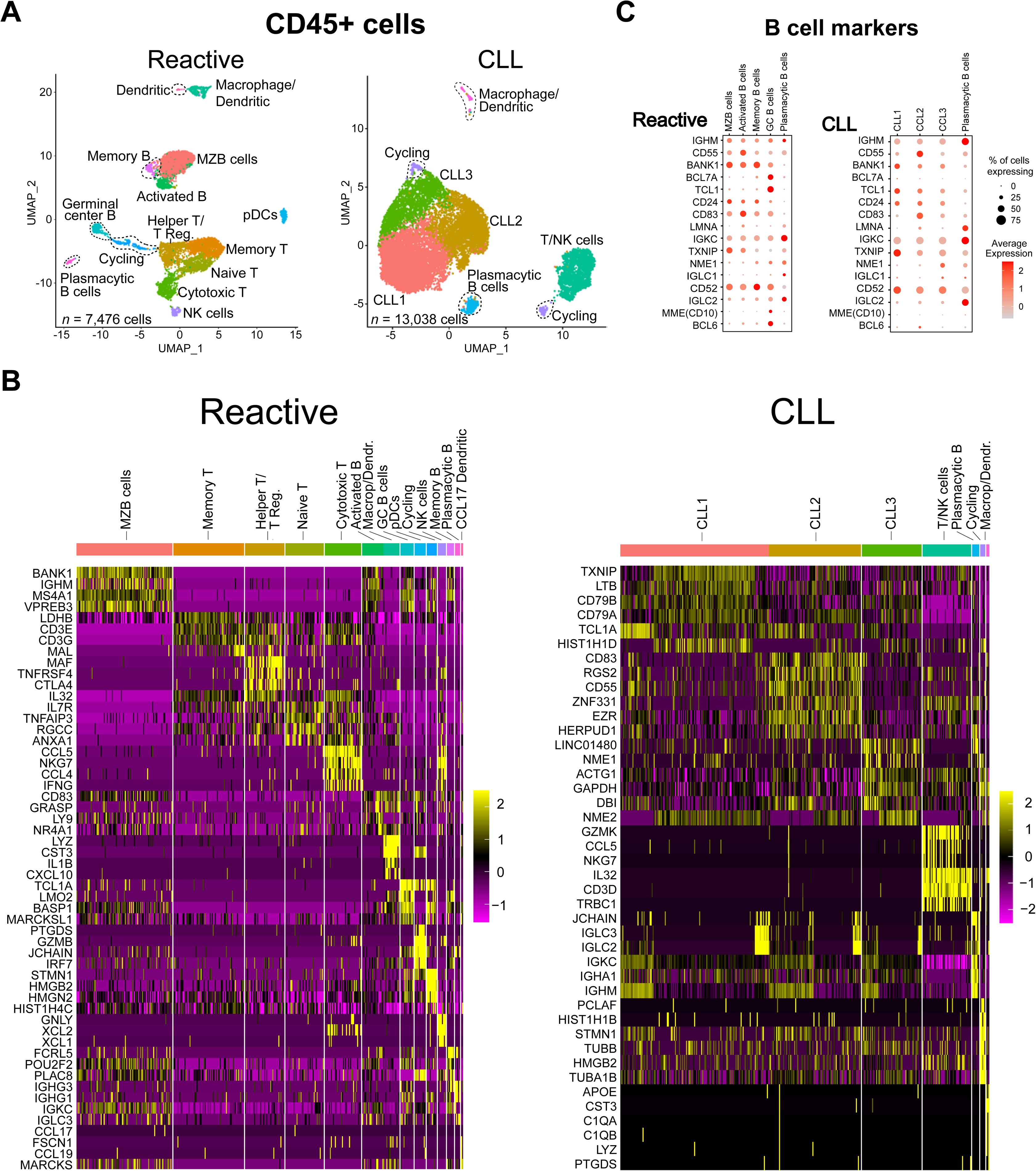
Major CD45^+^ cell types defined by scRNA-seq in LNs involved by reactive processes and CLL. **A)** UMAP plots corresponding to the integrated scRNA-seq datasets of of reactive and CLL LN CD45^+^ cells. **B)** Heatmaps showing top differentially expressed genes in the CD45^+^ cell fractions. **C)** Dot plots displaying the expression of top B cell genes in CD45^+^ cell clusters identified by scRNA-seq.

Additional analysis focused other immune cells in CLL and reactive LNs identified several T/NK cell and myeloid cell subsets and associated differentially expressed genes (Supplemental Figure 4, Supplemental Table 4). Prominent differences specific to CLL LNs included the presence of an expanded, well-defined cluster of *FOXP3*+ T regulatory cells (Tregs) and sharply decreased numbers of plasmacytoid dendritic cells (Supplemental Figure 4A and B, respectively), consistent with prior studies^30–32^.

### Expression of *MYC* and *MKI67* is Largely Discrete in Proliferating CLL Cells

Because proliferation of CLL cells is spatially confined to PCs, we next sought to determine how CLL cells transition through successive proliferative states within this niche. To address this question, we examined expression of MYC and Ki67 (encoded by MKI67), canonical markers of proliferation in B-cell lymphomas^33^, at single-cell resolution.

In our dataset, *MYC*^+^ and *MKI67*^+^ CLL cells mainly fell into two discrete UMAP clusters with little overlap (Figure 3A-C), with Slingshot trajectory analysis suggesting a precursor-product relationship (Figure 3A)^34^. Consistent with this idea, Seurat cell-cycle scoring indicated that *MYC*^+^ cells were enriched in G1/S phase cells, whereas *MKI67*^+^ cells were predominantly in G2/M phase (Figure 3D-E). Furthermore, *MCM6* and *PCNA*, genes that peak in expression in S phase, were most highly expressed in cells predicted by Seurat to be on a continuum between *MYC*^+^ and *MKI67*^+^ cells (Figure 3F). Finally, immunostaining showed that MYC^+^ and Ki67^+^ cells were largely non-overlapping within CLL PCs (Figure 3G-I). Together, these findings define an ordered proliferative program within PCs in which MYC is expressed early on following cell cycle entry and then declines as cells progress through the cell cycle and begin to express Ki67.

**Figure 3.**
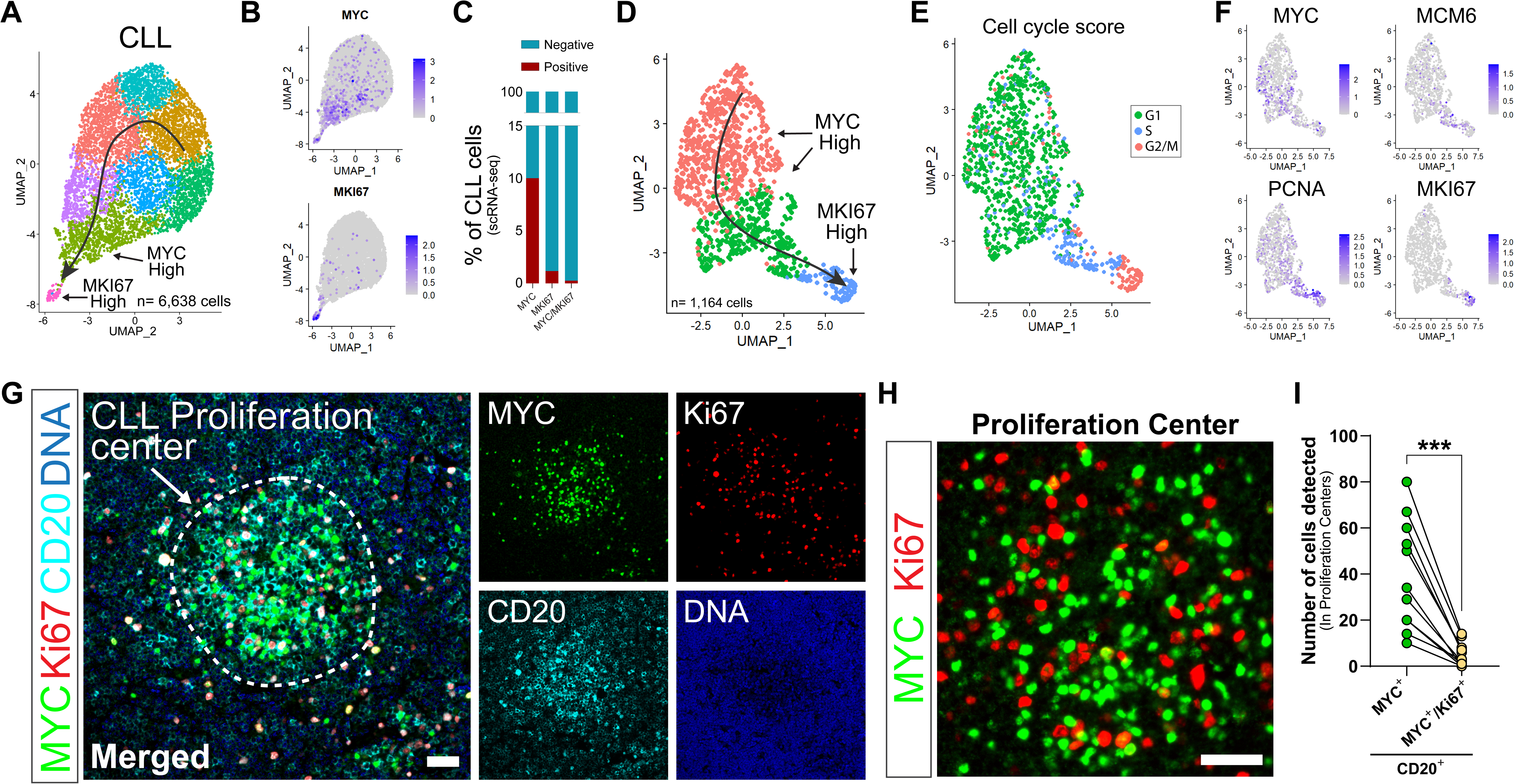
*MYC* and *MKI67* expression in CLL PCs is largely discrete. **A)** UMAP plot of CLL cells from specimen CLL1 with superimposed Slingshot trajectory analysis. **B)** UMAP feature plots showing *MYC* and *MKI67* expression in the subset of CLL cells from panel A. **C)** Percentage of CLL cells from an integrated scRNA-seq dataset (3 CLL patients) expressing *MYC*, *MKI67*, or both *MYC* and *MKI67*. **D)** UMAP plot showing *MYC*^+^ and *MKI67*^+^ CLL cells from panel A (subset of MYC and MKI67 clusters from 1A) with superimposed Slingshot trajectory analysis. **E)** Seurat cell cycle score showing CLL cells from panel D corresponding to cells in G1, S, and G2/M phases. **F)** Expression of selected cell cycle associated genes displayed on cells from panels D and E. **G)** Representative immunofluorescence microscopy images showing a CLL PC stained for MYC, Ki67 and CD20. Nuclei were counterstained with DAPI. Scale bar, 75 µm. **H)** Higher magnification image of the CLL proliferation center from G) showing MYC (green) and Ki67 (red) expression. Scale bar, 75 µm. **I)** Quantification of CD20⁺ cells expressing MYC alone or co-expressing MYC and Ki67 within 10 individual proliferation centers derived from 5 CLL-involved lymph nodes. ***P < 0.001, two-sided paired Wilcoxon signed-rank test.

### Characterization of CD45^-^ stromal cells in reactive and CLL LNs

The structural organization of normal and lymphoma-involved LNs is generated by cross talk between stromal cells such as fibroblasts and benign and neoplastic immune cells, and we therefore reasoned that tumor-specific features of CLL LNs such as PCs must stem from changes in the composition and distribution of resident CD45- stromal cells^35–37^. Integrated analysis of scRNA-seq of CD45^-^ cells from CLL and reactive LNs identified several types of fibroblasts and endothelial cells as well as plasma cells (Figure 4A), each defined by differentially expressed genes, as shown in heat maps (Figure 4B) and dot plots (Supplemental Figure 5A, B) (for gene lists, see Supplemental Table 5).

**Figure 4.**
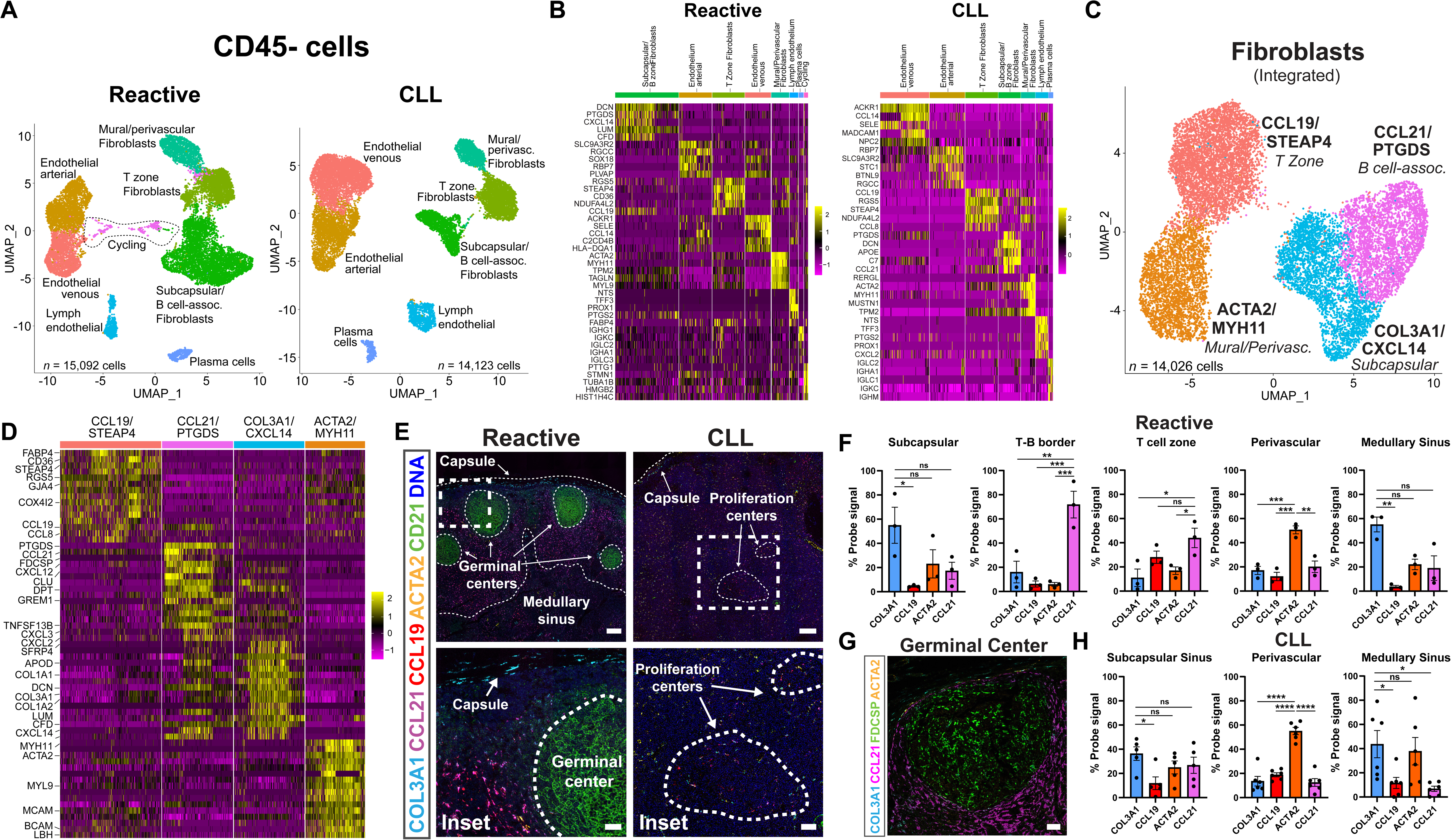
Identification and location of CD45^-^ stromal cell types in reactive and CLL LNs. **A)** UMAP plots displaying the integrated scRNA-seq datasets of CD45^-^ cells from LNs involved by reactive hyperplasia or CLL. **B)** Heatmaps showing expression of top differentially expressed genes in reactive and CLL CD45^-^ LN cells shown in A). **C)** UMAP plots displaying the integrated scRNA-seq datasets of all fibroblasts from reactive and CLL LNs. **D)** Heatmaps showing expression of top differentially expressed genes in the 4 main fibroblast types shown in C). **E)** Confocal immunofluorescence microscopy images of sections of LNs involved by reactive hyperplasia or CLL stained with RNAscope probes for *COL3A1*, *CCL21*, *CCL19*, and *ACTA2*. *COL3A1* and *CXCL14* probes were tagged with the same fluorophore to enhance detection of *COL3A1*/*CXCL14*-expressing fibroblasts. FDCs were detected by staining for CD21 protein. CLL PCs were spatially localized by immunostaining for MYC on an immediately adjacent serial section. Scale bars, 400 µm (top) and 50 µm (bottom, inset). **F)** Quantification of RNAscope probe signal for *COL3A1*/*CXCL14*, *CCL19*, *ACTA2*, and *CCL21* across anatomically defined regions of reactive lymph nodes, including subcapsular sinus, T–B border, T cell zone, perivascular regions, and medullary sinus. RNAscope signal is expressed as the percentage of total image area occupied by each probe. Each dot represents an individual lymph node sample; bars indicate mean ± SEM. Statistical comparisons were performed using one way ANOVA followed by Dunnett’s multiple comparisons test for normally distributed data, or the Kruskal–Wallis test followed by Dunn’s multiple comparisons test for non normally distributed data; ns, non-significant; *P < 0.05; **P < 0.01; ***P < 0.001. **G)** Confocal image of a reactive germinal center showing spatial organization of fibroblast-associated RNAscope signals for *COL3A1*/*CXCL14*, *FDCSP* and *ACTA2*. Scale bar, 50 µm. **H)** Quantification of RNAscope probe signal for *COL3A1*/*CXCL14*, *CCL19*, *ACTA2*, and *CCL21* across anatomically defined regions of CLL lymph nodes, including subcapsular sinus, perivascular regions, and medullary sinus. RNAscope signal is expressed as the percentage of total image area occupied by each probe. Each dot represents an individual lymph node sample; bars indicate mean ± SEM. Statistical comparisons were performed using one-way ANOVA; ns, non-significant; *P < 0.05; ****P < 0.0001.

Among stromal cells, fibroblasts are crucial for maintaining lymph node structure through the production of collagen and other extracellular matrix components, while also coordinating immune cell interactions^4^. Fibroblasts have also been linked to supporting cancer cell growth and survival^3^, including in B cell tumors such as follicular lymphoma, in which FDCs, fibrobast-like cells, underlie the follicular growth pattern of the tumor cells^38^. We therefore next focused our analysis on LN fibroblasts from reactive and CLL LNs, leading to the identification of 4 major fibroblast clusters (Figure 4C). Heatmaps of highly expressed genes (Figure 4D, Supplemental Table 5) and UMAP projections (Supplemental Figure 6A) showed that fibroblasts in these clusters were characterized by higher expression of *COL3A1*/*CXCL14*, *CCL21*/*PTGDS*, *CCL19*/*STEAP4*, and *ACTA2*/*MYH11*, respectively, and were present in both reactive and CLL LNs (Supplemental Figure 6B).

To map and compare the spatial distribution of fibroblasts expressing cluster-defining markers, we performed *in situ* hybridization (ISH) and immunostaining studies on reactive and CLL LNs (Figure 4E-H; see Supplemental Figure 6C for higher power images). In reactive LNs, which contain B-cell follicles or germinal centers surrounded by T-cell zones, *CCL19*^+^ fibroblasts localized to T-cell zones, consistent with these cells being T-cell zone FRCs^5^, while *CCL21*^+^ fibroblasts were present in subcapsular areas, T-cell areas, and around B-cell follicles, localization patterns specific for marginal reticular cells and T-B border fibroblasts^39^. By contrast, *COL3A1*^+^ fibroblasts mainly localized to subcapsular regions and medullary sinuses, whereas *ACTA2*^+^ fibroblasts were mainly found in perivascular areas, consistent with these cells being mural and perivascular fibroblasts. Thus, each of the 4 major fibroblast subtypes identified in our initial cluster analysis have distinct anatomic locations in reactive LNs.

Additional analysis of LN fibroblasts at higher resolution identified 7 fibroblasts clusters with distinct sets of differentially expressed genes (Supplemental Figure 6D-E) that appear to encompass the 7 main fibroblast subtypes previously defined in murine lymph nodes^4,5,39^, indicating substantial conservation of LN fibroblast subtypes between humans and mice. An important fibroblast subtype identified through this deeper analysis corresponds to follicular dendritc cells (FDCs), which were readily detectable by CD21 immunostaining (Figure 4E) or *in situ* staining for *FDCSP* (Figure 4G) in reactive LNs. By contrast to reactive LNs, in CLL LNs *CCL19*^+^ and *CCL21*^+^ fibroblasts were redistributed widely and CD21^+^ FDCs were entirely absent (Figure 4E; Supplemental Figure 6E). Thus, effacement of CLL LNs is reflected by substantial changes in the spatial distribution and differentiation states of nodal fibroblasts.

Fibroblasts are increasingly recognized as sources of factors that influence the behavior of benign and malignant immune cells^40^, and we therefore used dot plot visualization to compare expression of factors implicated in CLL pathobiology^41^ in CD45^+^ and CD45^-^ cells (Supplemental Figure 7A, B). We observed that subcapsular/B-cell related fibroblasts express high levels of *TNFSF13B* (BAFF), a crucial cytokine for B cell survival and maturation^42,43^, in all LNs analyzed, suggesting these cells are an important source of this factor. Macrophage/dendritic cells also expressed relatively high levels of *TNFSF13B* (Supplemental Figure 6A). By contrast, despite prior work showing that *CD40LG*, *IL21*, and *TLR9* can promote the *ex vivo* expansion of CLL cells^15^, we identified only low level expression of these genes in CLL LNs. Increased stroma-dependent Notch signaling has been linked to worse outcomes in CLL^44^, and our data also revealed a complex pattern of expression of Notch ligands and receptors in CLL cells, immune cells, and stromal cells. Notably, several Notch ligands are highly expressed by CD45^−^ stromal cells in CLL LNs, in line with observations showing that the LN microenvironment enhances Notch activation in CLL cells ^45^.

As noted in Figure 1F, CLL LNs contained larger numbers of endothelial cells than reactive LNs, and we therefore performed a deeper analysis of LN endothelium. Integrated analysis identified clusters corresponding to blood endothelium (Supplemental Figure 8A-E) as well as high endothelial venule endothelial cells (Supplemental Figure 8F, G; see Supplemental Table 5 for differentially expressed genes). Relative to reactive LNs, CLL LNs yielded larger numbers of vein endothelial cells and smaller numbers of arterial endothelial cells, whereas high endothelial venule endothelial cells, identified by expression of *CHST4* together with *SELE*, *ICAM1*, and *VCAM1^46^*, were recovered in similar numbers (Supplemental Figure 8E, F). Cells corresponding to all major lymphatic endothelium cells types^47^ were also recovered from both reactive and CLL LNs (Supplemental Figure 8H-J), with CLL LNs yielding larger numbers of collecting, medullary, and subcapsular ceiling cells and fewer subcapsular floor lymphoendothelial cells (Supplemental Figure 8J).

### *CCL19*^+^ fibroblasts selectively localize to proliferation centers in CLL LNs

The PC, a histopathologic hallmark of CLL that is readily recognized by light microscopy, is defined by the presence of loose clusters of enlarged, proliferating CLL cells that are MYC^+^.^13^ Given the importance of fibroblasts in supporting LN architecture, we reasoned that this unique niche might be associated with a specialized fibroblast population. To explore this idea, we performed *in situ* studies to determine the spatial relationship between fibroblasts expressing cluster-defining markers and CD20^+^MYC^+^ PC CLL cells. We identified a population of *CCL19*-high/*CCL21*-low fibroblasts, which we named CLL proliferation-associated (CPA) fibroblasts, that is significantly enriched in PCs as compared to fibroblasts expressing other subtype-specific genes (Figure 5A-B). We also observed that some fibroblasts at the periphery of PCs co-express *CCL19* and *CCL21*, suggesting a border region containing fibroblasts transitioning between *CCL21*-high and *CCL19*-high cell states (Figure 5A-B, Supplemental Figure 9A).

**Figure 5.**
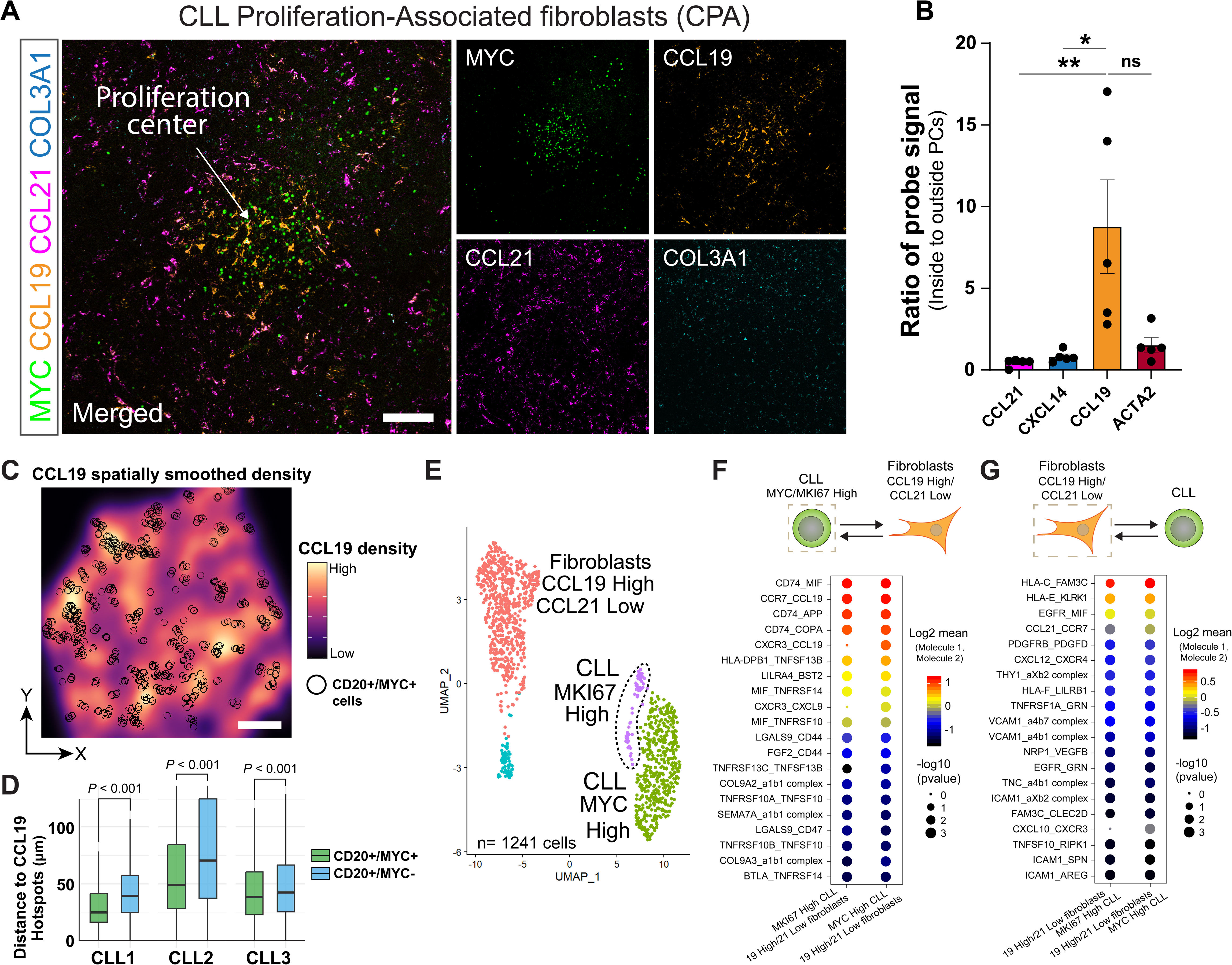
Identification, predicted interactions, and spatial transcriptomics of PC-associated fibroblasts in CLL LNs. **A)** Confocal images of dual immunofluorescence staining for MYC protein and *CCL19*, *CCL21*, *COL3A1*/*CXCL14* transcripts in a CLL PC. *COL3A1* and *CXCL14* probes were tagged with the same fluorophore to enhance detection of *COL3A1*/*CXCL14* expressing fibroblasts. Scale bar, 100 µm. **B)** Quantification of the hybridization signals detected inside and outside PCs in 5 LNs involved by CLL. Each dot represents an individual lymph node sample; Bars indicate mean ± SEM. Statistical comparisons were performed using the Kruskal-Wallis test followed by Dunn’s multiple comparisons test. *ns*, non-significant; \**P* < 0.05; \*\**P* < 0.01 **C)** Spatial transcriptomics of a CLL LN showing *CCL19* expression as smooth density with clustered *MS4A1*^+^*MYC*^+^ cells overlaid. Cells were slightly enlarged for visualization purposes. Scale bar, 400 µm. **D)** Average distances of clustered *MS4A1*^+^MYC^+^ and *MS4A1*^+^MYC^-^ cells to *CCL19*high “hotspots” (top 1% of signal) in 3 CLL LNs. Statistical comparisons were performed within each CLL case using two-sided Wilcoxon rank sum tests. **E)** UMAP plot of *MKI67-High* and *MYC-High* CLL cells and *CCL19-High/CCL21-Low* fibroblasts (predicted CPAs) (see Supplemental Materials for filtering steps). **F-G)** Top 20 ligand-receptor interactions between *MKI67/MYC*-High CLL cells and *CCL19-*High/*CCL21-*Low fibroblasts as predicted by CellPhoneDB. Circle size represents the -Log10 of P value for a specific interaction; thus, the larger the circle size the smaller the P value. The color of circles indicates expression levels of interacting molecules. On the left (F), CLL cells are the “anchor” for the analysis, whereas on the right (G) the *CCL19-*High/*CCL21-*Low fibroblasts are the “anchor”, indicated by a dashed square and dashed rectangle, respectively.

To further confirm the localization of cells expressing these markers in PCs, we performed Xenium spatial transcriptomic analysis on three CLL LNs with typical morphologic features, generating a combined dataset of 790,965 cells across the three profiled samples. Nearest-neighbor spatial analysis showed that *MS4A1*^+^(CD20)*MYC*⁺ cells were positioned closer to *CCL19*-expressing cells than were *MS4A1*^+^(CD20)*MYC*⁻ cells (Figure 5C-D), in agreement with the *in situ* staining results.

To identify additional markers that define CPA fibroblasts we integrated the scRNA-seq data from 3 CLL patient samples, focusing on fibroblast populations with high *CCL19* expression and low *CCL21* expression (see supplemental methods), allowing us to identify differentially expressed genes. In addition to *CCL19*, the most highly differentially expressed genes in this population include *CALD1*, *CST3*, and *CPE*, genes linked to chemotaxis, cytoskeletal organization, inflammatory programs, and peptide processing, respectively (Figure 5E, Supplemental Figure 9B-C).

We next used CellPhoneDB, a bioinformatic tool that integrates scRNA-seq data with curated ligand-receptor pairs^48^, to predict likely ligand-receptor interactions involving *CCL19*-high/*CCL21*-low fibroblasts and *MYC* or *MKI67*-high CLL cells based on genes that are highly expressed in these cell populations (Figure 5E). The top predicted interacting ligands and receptors included CD74 (which is widely expressed in B cell lymphomas^49^) and its ligands MIF (macrophage migration inhibitory factor), COPA, and APP (Figure 5F-G, Supplemental Table 6). Best known as a chaperone for HLA class II molecules, CD74 interacts with receptors such as CXCR4 to facilitate MIF-induced ERK/MAP kinase and PI3K/AKT signaling^50^, pathways involved in proliferation and survival. Another top predicted interaction was CCL19 and CCR7. CCL19 is linked to chemotaxis of CCR7^+^ cells and activates survival signaling through pathways such as PI3K/AKT and MAPK/ERK. In CLL LNs, *CCL19* expression is restricted to fibroblasts while *CCR7* is expressed by CLL cells (as well as T cells, Supplementary Figure 10A), suggesting that CCL19^+^ fibroblasts may help recruit or retain CCR7^+^ PC CLL cells. This possibility is consistent with our spatial transcriptomics data, which showed that *MS4A1*^+^(CD20)*MYC*^+^*CCR7*⁺ CLL cells were closer to *CCL19*-rich areas than were *MS4A1*^+^(CD20)*MYC*⁻ CLL cells (Supplementary Figure 10B-C), as well as immunostaining results, which confirmed that CCR7 is expressed within proliferating CLL cells in PCs (Supplementary Figure 10D).

### Predicted ligand-receptor pairs between proliferating CLL cells and immune cells

Because MYC expression in cancer cells has been linked to the establishment of an immunosuppressive microenvironment^51^, we next sought to determine whether proliferating *MYC*⁺ or *MKI67*⁺ CLL cells engage in distinct interactions with immune cells within the LN microenvironment. We first investigated which types of immune cells are likely to preferentially interact with proliferating CLL cells using CellChat^52^ and CellPhoneDB. The highest number of predicted interactions between proliferating CLL cells and immune cells involved macrophage/dendritic cell populations, followed by cytotoxic T/NK cells (Figure 6A-C, and Supplemental Table 7), consistent with immunostaining results in 13 CLL LNs, which confirmed the presence of CD68^+^ macrophages and CD8^+^ cytotoxic T cells near MYC^+^ CLL cells in PCs (Supplemental Figure 11A- C).

**Figure 6.**
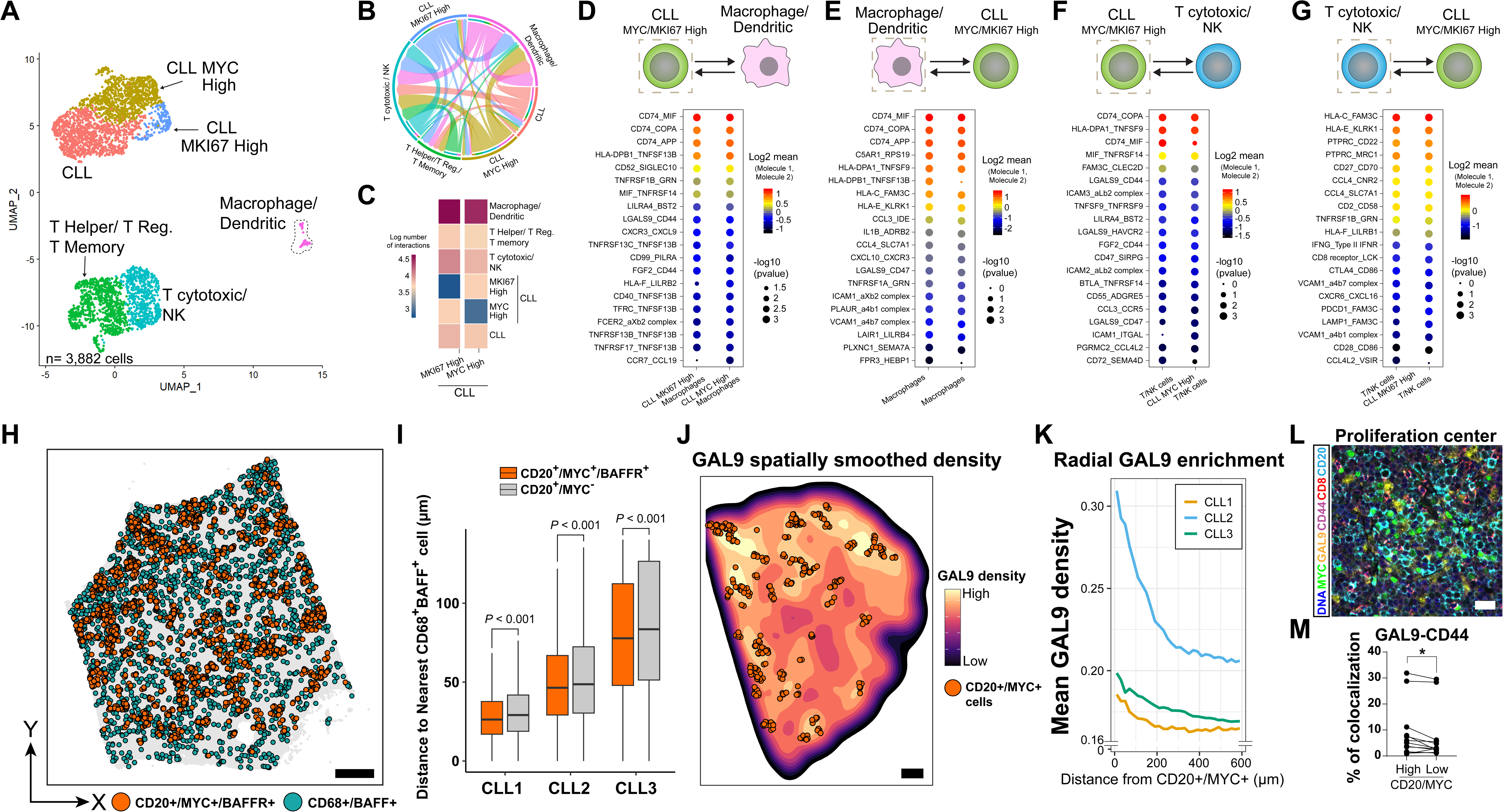
Predicted ligand-receptor interactions between CLL cells and immune cells. **A)** UMAP plots showing *MKI67*^+^ and *MYC*^+^ CLL cells and T cells, NK cells, macrophages, and dendritic cells. **B)** Chord diagram showing CellChat-predicted interactions between MKI67-high and MYC-high CLL cells and immune cell populations. **C)** Heatmaps showing the number of CellPhoneDB-predicted interactions between MKI67-high and MYC-high CLL cells and immune cell populations. **D-G)** Top 20 CellPhoneDB-predicted ligand-receptor interactions between *MKI67*^+^ and *MYC*^+^ CLL cells and macrophage/dendritic cells (D-E) and T cytotoxic/NK cells (F-G). Cells on the left serve as the “anchor” for the analysis, indicated by a dashed squares. Circle size represents the -Log_10_ of P value for a specific interaction (larger circles = higher significance). Circle color indicates expression levels of interacting molecules. **H)** Spatial distribution of *MS4A1*^+^*MYC*^+^*TNFRSF13C*⁺(BAFFR) cells and *CD68*⁺*TNFSF13B*⁺(BAFF) macrophages in a CLL LN. Cells were slightly enlarged for visualization purposes. Scale, 400 µm. **I)** Nearest-neighbor analysis comparing distances between *MS4A1*^+^*MYC*^+^*TNFRSF13C*⁺ and *MS4A1*^+^MYC⁻*TNFRSF13C*⁺ CLL cells and the nearest *CD68*⁺*TNFSF13B*⁺ macrophage in three different CLL LNs. Statistical comparisons were performed within each CLL case using two-sided Wilcoxon rank sum tests. **J)** Spatial transcriptomics showing smoothed spatial estimates of *LGALS9* expression overlaid with clustered *MS4A1*^+^*MYC*^+^ CLL PC cells (orange). Color scale represents normalized *LGALS9* expression density. Cells were slightly enlarged for visualization purposes. Scale bar, 400 μm. **K)** Mean *LGALS9* expression plotted as a function of distance from clustered *MS4A1⁺MYC⁺* CLL PC cells. **L)** Immunofluorescence confocal image of CLL LN showing staining for GAL9, MYC, CD20, CD44, CD8, and DNA within a CLL PC. Scale bar, 25μm. **M)** Quantification of GAL9-CD44 colocalization near CD20^+^MYC^+^ and CD20^+^MYC^-^ CLL cells. (N = 13 cases). Each pair of points represents one case. *P < 0.05, two-sided paired Wilcoxon signed rank test.

We next used CellPhoneDB to identify specific ligand-receptor pairs involving these cells. These pairs can sorted into 4 functional categories: i) progrowth and survival (e.g., TRNFRSF13C-TNFSF13B, CD74-MIF; ii) cell motility (e.g., CD55-ADGRE5)^53^; iii) immunoevasion (e.g, HLA-F-LILRB2, GAL9-HAVCR2 (TIM3), GAL9-CD44, CD86-CTLA4)^54–56,57^; and iv) cell-cell adhesion (e.g., ICAM1-aXβ2, ICAM3-aLβ2 (Figure 6D-G). Importantly, most of the top ligand-receptor pairs predicted by CellPhoneDB were also identified by NicheNet (Supplemental Figure 12A-C), another bioinformatic tool that predicts ligand-receptor pair interactions based on signaling pathway activity in receiver cells inferred from RNA expression^58^.

To begin to evaluate the anatomic distribution of top predicted ligand-receptor pairs in the CLL proliferative niche, we used spatial transcriptomics (see Supplemental Table 7 for gene sets). These analyses showed that macrophages expressing *TNFSF13B* (BAFF) and *ICAM1* were enriched near CLL cells in PCs expressing their cognate receptors, *TNFRSF13C* (BAFFR) and components of the integrin αXβ2 heterodimer (*ITGAX*-*ITGB2*), respectively (Figure 6H-I, Supplemental Figure 13A-B). Spatial transcriptomic data also showed that *LGALS9* (Galectin 9 or GAL9) expression, a gene encoding a secreted protein previously linked to cytotoxic T cell immune suppression ^54^, was elevated near CLL cells in PCs (Figure 6J-K, Supplemental Figure 13C), suggesting that PCs constitute an immunosuppressive niche. To further evaluate predicted ligand-receptor interactions, we performed multiplex confocal immunofluorescence microscopy in 10 additional CLL LN cases (n=13), focusing on two of the top ranked interactions from our bioinformatic analyses, GAL9-CD44 and CD74-MIF. GAL9-CD44 colocalization (Supplemental Figure 13D) was detected in all cases examined. Because GAL9 is predicted to be secreted by MYC⁺ CLL cells, we quantified its spatial distribution and found higher GAL9 signal within a 30 µm radius^59^ of CD20⁺MYC⁺ cells compared with regions outside of PCs without CD20^+^MYC^+^ cells (Figure 6L-M, Supplemental Figure 13D), as predicted from the spatial transcriptomic results.

### CD74 signaling promotes CLL growth

Similar in situ analyses of CD74 and MIF showed strong colocalization of CD74 and MIF on CD74^+^ CLL cells (Figure 7A-B). Unlike GAL9, there was no enrichment of MIF and CD74 colocalization near MYC^+^ CLL cells (Figure 7B), possibly due to expression of these molecules in multiple cell types (Figure 7C-D) and secretion of MIF into the local microenvironment. To study the function of CD74 signaling, among the top pathways predicted to be active in CLL cells, we established a CLL spheroid culture system modified from a recently described method^60^. Under optimized culture conditions, primary CLL cells form compact spheroids and actively divide. Growth of both IGHV-mutated and IGHV-unmutated CLL cells was significantly inhibited by a monoclonal antibody specific for CD74, whereas an isotype control antibody had no effect (Figure 7E-G, and Supplemental Figure 14). Given the consistent expression of CD74 by CLL cells and widespread expression of CD74 ligands in the CLL microenvironment, these findings support a role for CD74 signaling in promoting CLL growth *in vivo*.

**Figure 7.**
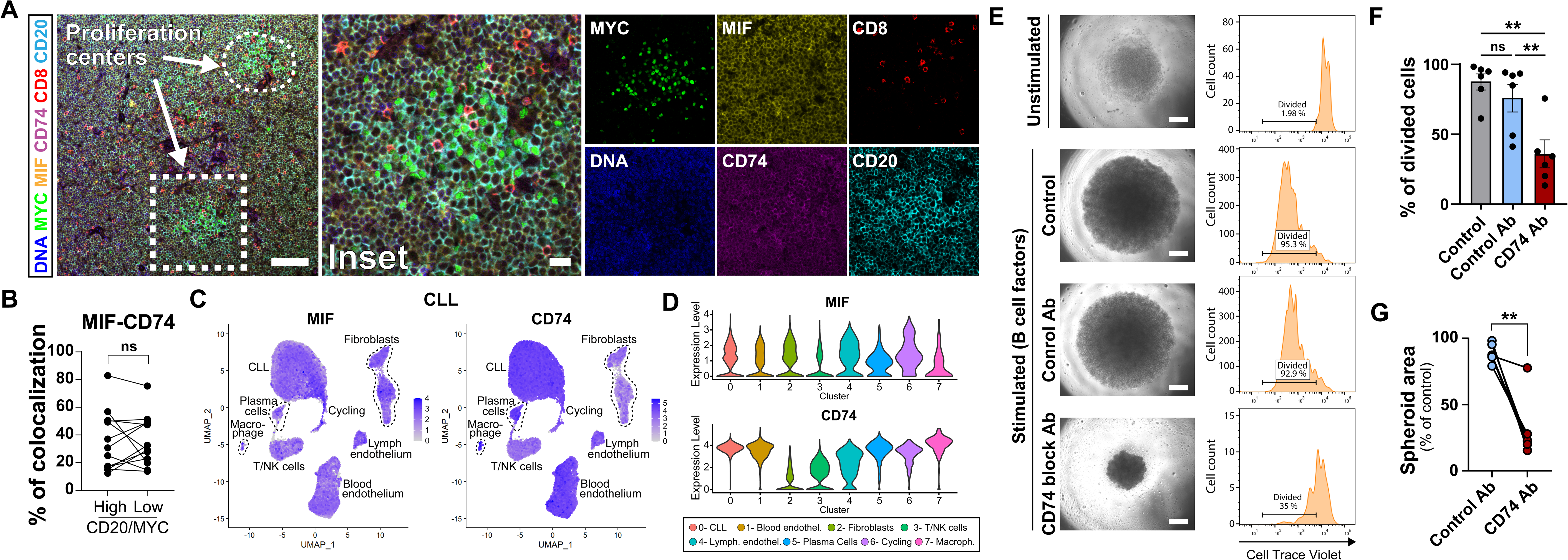
Nodal expression and functional relevance of CD74 and MIF in CLL. **A)** Immunofluorescence confocal microscopy images depicting staining of CD74 and MIF within CLL PCs. Inset signals are shown individually in the right-hand panel. Scale bars, 100 μm (left), 25 μm (right, inset). **B)** Quantification of colocalization of CD74-MIF within CLL PCs on CD20^+^MYC^+^ cells (N = 13 cases). Each pair of points represents one case, with circles indicating the average of all detected signals per case. MYC^+^ and MYC^-^ areas were compared using a two-sided paired Wilcoxon signed rank test. ns: non-significant. **C)** UMAP feature plots showing expression of MIF and CD74 across integrated scRNA-seq datasets from 3 patient CLL lymph nodes. **D)** Violin plots showing MIF and CD74 expression levels across the major cell populations identified in the integrated scRNA-seq datasets, including CLL cells, blood endothelial cells, fibroblasts, T/NK cells, lymphatic endothelial cells, plasma cells, cycling cells, and macrophages. **E)** Growth of CLL cells in spheroid cultures. Left, representative brightfield images; right, cell proliferation assessed by CellTrace Violet dilution over 7 days. Representative results are shown for CLL cells incubated in the absence (unstimulated) or presence of added factors, a CD74 blocking antibody, or an isotype control antibody. Scale bars, 100 µm. **F-G)** Quantification of CLL cell growth as assessed by cell division **(F)** or spheroid size **(G)** after 7 days in culture under the indicated conditions. Each dot represents an independent CLL case (N = 6). Statistical comparisons were performed using one-way ANOVA followed by Dunnett’s multiple comparisons test in (F) and two sided Wilcoxon rank sum test in (G). NS, non-significant; **P < 0.01.

## Discussion

Building on a generally applicable LN pipeline, we used scRNA-seq, bioinformatics, spatial transcriptomics, microscopy-based, and functional ex vivo culture approaches to characterize proliferating CLL cells in LNs and gain insight into the factors that support their growth, survival, and immune evasion.

Our work indicates that *MYC* expression is transient in proliferating CLL cells, falling as cells accumulate *MKI67* transcripts. This is in line with studies in normal germinal center B cells showing that Ki67 levels are low in G1 and peak in G2/M^61^, whereas MYC levels rise quickly in G1 phase and then fall during cell cycle progression^62,63^. Notably, in germinal center B cells MYC abundance appears to govern proliferative output, such that the magnitude of MYC induction limits the number of subsequent cell divisions^64^. Although LNs involved by CLL lack germinal centers, our findings suggest proliferating CLL cells engage analogous regulatory circuits that drives limited, microenvironment-dependent MYC upregulation. Of interest, acquired genetic lesions during CLL progression that dysregulate MYC are associated with Richter transformation^65^, highlighting the need to elucidate mechanisms leading to *MYC* downregulation during cell cycle progression^64,66^.

In line with the striking architectural differences in reactive LNs and CLL LNs, we observed profound changes in nodal CD45^-^ cells in CLL LNs, including redistribution of *CCL19*^+^ and *CCL21*^+^ fibroblasts, increased vascularity, and depletion of follicular dendritic cells. Of particular interest, we find PCs are enriched for a distinct population of *CCL19*-high/*CCL21*-low fibroblasts (termed “CPA fibroblasts”) that appears to be a rich source of factors implicated in CLL cell trafficking, survival, and microenvironmental support, and that may recruit CLL cells to PCs through a CCL19-CCR7 signaling axis. This axis has been implicated previously in CLL lymph node homing and retention^67,68^, and our findings extend this association by suggesting that specialized CCL19^+^ fibroblasts within PCs may contribute to organization of the proliferative niche. Interestingly, *CCL19*^+^ fibroblasts have recently been linked to tertiary lymphoid organ formation in colorectal cancer liver metastasis by recruiting CCR7^+^ B cells^69^. Further work aimed at functional characterization of CPA fibroblasts will be important in defining their contribution to the pathobiology of CLL.

Using interactions initially predicted through informatics approaches, we also validated the colocalization of the ligand pairs GAL9-CD44 and CD74-MIF. We find that GAL9-CD44 colocalization is greatest near MYC^+^ CLL cells in PCs, suggesting MYC-expressing cells enhance GAL9-dependent local immunosuppression, consistent with other work linking MYC to immunosuppression^51,70^. This is also in line with our observation that cytotoxic CD8⁺ T cells near MYC^+^ CLL cells are in a microenvironment containing higher levels of GAL9, an association consistent with a recent study suggesting that T regs and exhausted T-cell states in CLL are modulated by GAL9^71^. Our analyses also reveal that MYC^+^ CLL cells reside in close physical proximity to BAFF^+^/ICAM1^+^ macrophages, consistent with a supportive role for macrophages in providing survival and adhesion signals to proliferating CLL cells. Altogether, these findings suggests that PCs represent a progrowth, proadhesion, immunoprotective niche.

CD74–MIF signaling has previously been implicated in the growth and survival of multiple cancers, including CLL in studies using 2D culture systems and mouse models^72–75^. Here, building on transcriptomic and spatial analyses of CLL LNs and using a new 3D spheroid model that consistently allows for expansion of primary human CLL cells *ex vivo*, we provide additional evidence supporting the importance of CD74 signaling in CLL growth. Of note, CD74 signaling has also been associated with resistance to immune checkpoint therapy in multiple cancers^76^. Inhibitors targetting MIF, one of several CD74 ligands, are in clinical development,^77^ and our work provides an additional rationale for further evaluation of CD74 signalling as a therapeutic target in CLL.

## Acknowledgements

The authors thank the Harvard Cancer Consortium Specialized Histopathology Core for IHC staining technical assistance, in particular Caitlin Edwards and Teresa Bowman. scRNA-seq and Xenium spatial transcriptmoics were performed by the Center for Cellular Profiling at Brigham and Women’s Hospital; we are thankful to Kevin Wei, the Center director, for insightful thoughts at the earlier stages of the project, as well as Junning Case, Tamara Salloum and Sean Prell for technical support. We thank the team of the Microscopy Resources on the North Quad (MicRoN) of Harvard Medical School, particularly Paula Montero Llopis, the Resource director, Praju Anekal, Adrienne Wells, and Ryan Stephansky for their insightful thoughts on the microscopy panels; Kelly Street (University of Southern California) for helpful discussion on Slingshot trajectory analysis; and Marco Haselager and Eric Eldering (University of Amsterdam) for advice on spheroid cultures. A.P.F is supported by a Lymphoma Research Foundation postdoctoral fellowship and the 2024 Cotran-Gimbrone Research Award. This work was also supported in part by R35 CA220340 (to S.C.B.), a microgrant from the Brigham Research Institute, a Doris Duke Charitable Foundation’s Physician Scientist Fellowship and American Society of Transplantation and Cellular Therapy New Investigator Award (to J.D.B.), and R01 AI091627 (to I.M.).

## Authorship and Conflict of Interest

A.P.F., S.W., S.S., M.D., D.D. and B.S. participated in biopsy collection. A.P.F. processed human lymph nodes and prepared samples for sorting and subsequent scRNA-seq. A.P.F. and D.M. performed in vitro functional assays. S.W. and A.P.F performed microscopy staining of FPPE patient samples. S.M.F. and J. Brown characterized and provided CLL peripheral blood mononuclear cell samples. A.P.F. developed and implemented the scRNA-seq/Seurat analysis pipeline. A.P.F. performed CellChat and trajectory analyses. A.P.F, L.P and M.E.L. performed CellPhoneDB analyses, A.P.F. and L. Poluben performed NicheNet and InferCNV analyses. A.P.F. performed the spatial transcriptomics data analysis. A.P.F and J.C.A. developed the N_Mito mitochondrial filtering algorithm. A.P.F. led all data analyses and prepared all figures and tables. J. Brandstadter, E.P. and I.M. provided conceptual advice and feedback on lymph node sample processing and tissue cryopreservation. J. Brown provided conceptual advice and feedback on in vitro functional assays. S.W., L. Pan, J.Brandstadter, E.P., D.M.D., B.S., M.E.L., S.R.S., J.Brown, I.M., and S.C.B. provided conceptual advice and feedback. All authors discussed the results and commented on the manuscript. A.P.F, S.C.B. and J.C.A. conceived the study. A.P.F. and J.C.A. wrote the manuscript with input from all the other authors. I.M. has received research funding from Genentech and Regeneron and is a member of Garuda Therapeutics’ scientific advisory board (all unrelated to this work). S.C.B. is on the board of directors for the non-profit Institute for Protein Innovation, is on the scientific advisory board and receives funding from Erasca, Inc., for an unrelated project; is an advisor to MPM Capital; and is a consultant for IFM, Scorpion Therapeutics, Odyssey Therapeutics, and Ayala Pharmaceuticals for unrelated projects. J.C.A. is a consultant for Ayala Pharmaceuticals and Remix Therapeutics and is on the scientific advisory board of Cellestia, Inc., all for work unrelated to that described herein.

## Methods

### Use of human tissues

LN biopsies were collected from patients with lymphadenopathy under IRB-approved protocol 14-076. Peripheral blood CLL cells were collected under IRB-approved protocol 01-206. Characteristics of the cases used are provided in Supplemental Table 1.

### LN collection and storage

Fresh LNs were cut into 1mm^3^ fragments, viably frozen in Cryostor® CS10 (Stem Cell Technologies), and stored at -80°C. Formalin-fixed paraffin-embedded (FFPE) LNs involved by CLL were from the archives of the Department of Pathology at Brigham and Women’s Hospital. Samples involved by reactive processes or CLL were selected after review by a hematopathologist (JCA). Diagnoses were per the World Health Organization Classification, 5^th^ edition^78^. Pathologic findings are provided in Supplemental Table 1.

### Sample processing for scRNA-seq

Cryopreserved lymph node samples were thawed and digested in Dulbecco’s Modified Eagle Medium (DMEM) containing 2% fetal bovine serum, 1mM CaCl_2_, 1mg/mL collagenase IV (Worthington Biosciences, cat. #LS004210), 0.6mg/mL collagenase P (Milipore Sigma, cat. #11213857001), 0.2 mg/ml liberase (Sigma, cat. # 05401020001), and 0.3 mg/ml DNAse I (Sigma, cat. #07469) at 37° C with stirring. Dispase II (0.4 mg/mL, cat. # D4693-1G) was added in cases in which the tissue was resistant to digestion. Every 10 min, the samples were mechanically disrupted by pipetting up and down with a Pasteur pipette, the supernatant was transferred to a new tube on ice, and additional digestion mix was added to non-digested tissue. This process was repeated for up to 1 hr. Cell clumps and debris in the supernatant were removed by passage through a mesh with 100 μm pores and red cells were lysed using red cell lysis buffer (Sigma, cat. #420301). Cells in suspension were then incubated for 10 min with Fc Blocking Reagent (Biolegend, cat. #422302) followed by DAPI (Biolegend, 1 µg/mL, cat. #422801) in FACS buffer (phosphate buffered saline [PBS] containing 2% fetal bovine serum). Cells were subsequently incubated with CD45-PE-Cy7 antibody (Biolegend, cat. #304015) for 30 min on ice. After washing 3 times in FACS buffer, cells were loaded into FACS tubes with a 40 µm mesh cap and sorted into CD45^+^ or CD45^-^ DAPI^-^ fractions on a 4-Laser BD FACSaria Fusion Cell Sorter at 4°C. Flow results were analysed with FlowJo software v10.7.2. Sorted cells were resuspended in 0.4% BSA in PBS at 1000 cells/µl. For all cases except reactive LN3, and CLL LN3, CD45^+^ and CD45^-^ cells were loaded independently on 10X Genomics Chromium chips. For reactive LN3, and CLL LN3, sorted CD45^+^ and CD45^-^ cells were mixed at a ratio of 3:7 and run on a single 10X Genomics Chromium chip.

### scRNA-seq

Encapsulation and cDNA library generation (10X Genomics Single Cell 3′Kit V3.1) from sorted cells were per the manufacturer’s protocol. cDNA libraries were sequenced to an average of 44,171 reads per cell using an Illumina Nextseq 500. scRNA-seq reads were processed with Cell Ranger v3.1 and aligned to the GRCh38 transcriptome with STAR aligner.

### scRNA-seq data processing and filtering

scRNA data were analysed with Seurat single cell analysis pipeline (version 4.1.1) using R (version 4.2.0). Trajectory analysis was done using Slingshot (version 2.4.0). Ligand-receptor interaction analysis was done using CellPhoneDB (version 2.1), CellChat (version 1.6.1) and NicheNet (version 2.0.0). Copy number variations were determined using InferCNV (version 1.16.0).

### scRNA-seq data processing and quality control filtering

The Seurat single cell analysis pipeline (version 4.1.1) and R (version 4.2.0) were used to analyze scRNA-seq data. Low quality cells expressing <200 genes were removed as were cell doublets, which were identified by i) gene counts above twice the average for CD45^+^ or CD45^-^ cell populations (nFeature_RNA) or (ii) co-expression of genes belonging to cells of different lineages (e.g., B and T cell-specific genes). To retain healthy cells with high numbers of mitochondrial transcripts, we created the filter (N_Mito), which is based on the ratio of mitochondrion genome-encoded transcripts (M_MT) to nuclear transcripts encoding mitochondrial proteins (N_MT) or selected housekeeping gene transcripts (*GAPDH*, *ACTB*, *PGK1*, and *B2M*). Nuclear genes encoding mitochondrial proteins were derived from a public, curated list of 1136 genes (MitoCarta 3.0, Broad Institute). From this list, we selected 81 genes (Supplemental Table 2) that were ubiquitously expressed in all identified UMAP-defined clusters. The metric N_Mito is defined as N_Mito = ∑%MT_genes / ∑(%N_MT + %housekeeping genes). Retained cells had an N_Mito “score” of less than 7.5. The rationale for retaining cells with an N_Mito score of <7.5 was based on the histogram distribution of scores, which showed an inflection point at this value. To maximize the number of cells retained for downstream analyses, we also retained cells with an MT score of <15%, a threshold that also corresponded to an inflection point in the distribution of scores for this metric. After filtering, data analysis was performed with the Seurat package functions NormalizeData, FindVariableFeatures, ScaleData, RunPCA, FindNeighbours, FindClusters, and RunUMAP. In brief, gene expression data were normalized by total expression, multiplied by a scale factor of 10,000, and log transformed. The 2,000 most variably expressed genes were identified and expression of these genes was scaled before performing principal component analysis. Twenty principal components were used for graph-based clustering and UMAP dimensionality reduction. Integration of data from different samples was performed using Harmony (version 0.1.0). Trajectory analysis was performed using Slingshot (version 2.4.0). Ligand-receptor interactions between cells were inferred using CellPhoneDB (version 2.1). The top 20 pairs of cognate ligands and receptors, defined as the pairs with the lowest P values and highest gene expression, were selected and ranked based on expression levels. CellChat (version 1.6.1) was additionally used to calculate total number of cell-cell interactions betwemm CLL cells and immune cells. NicheNet (version 2.0.0) was also used to infer ligand-receptor pairs based on their downstream effects on target gene expression networks in receiver cells, using reactive lymph nodes as a reference state. Cells of similar lineage were used as corresponding controls in the reactive samples. InferCNV (version 1.16.0) was used to infer copy number variations (CNVs) in CLL cells and plasmacytic cells in CLL case 1, using T helper cells as reference euploid cells.To determine CPA identity, scRNA-seq data from three integrated CLL patient samples were further subsetted to identify CCL19-high/CCL21-low fibroblasts, defined as fibroblasts with CCL19 expression >2 and CCL21 expression <2. Similarly, MYC-high and MKI67-high CLL cell populations were defined as cells with MYC or MKI67 expression >0.5 within the same integrated dataset. Differential gene expression and ligand-receptor interaction analyses were then performed on these populations to infer transcriptional programs and candidate cellular interactions associated with proliferative niches.

### Formalin-fixed paraffin-embedded section preparation

Formalin-fixed paraffin-embedded lymph nodes were sectioned at 5 µm with a Leica RM2255 or a Thermo Scientific Shandon Finesse ME microtome using RNase-free technique and applied to Superfrost+ charged glass slides for downstream staining reactions.

### *In situ* hybridization

*In situ* hybridization was performed using the manual RNA Scope Multiplex Fluorescent V2 kit (ACD-Bio) according to the manufacturer’s protocol. FFPE tissues were cut at 5 microns. *In situ* hybridization was done using the RNA Scope Multiplex Fluorescent KitV2 (ACDbio, #323120) using probes conjugated to Opal dyes. Immunostaining on FFPE sections was performed with HRP-based tyramide signal amplification. RNAscope probes obtained from ACD-Bio for human *CXCL14* (554648-C2), *COL3A1* (549431-C2), *CCL21* (N878391-C3), *FDCSP* (444231-C3), *CCL19* (474361), or *ACTA2* (311811-C4) were conjugated to OPAL color dyes (AKOYA Biosciences) to permit visualization.

### Fibroblast subtype identification in tissue sections

Because *CCL19* was expressed more consistently than *STEAP4* in cells of the *CCL19*/*STEAP4* cluster and *PTGDS* expression was not confined to fibroblasts, we used the combination of *CCL21*-high/*CCL19*-low staining to identify *CCL21*/*PTGDS* fibroblasts and *CCL19*-high/*CCL21*-low staining to identify *CCL19*/*STEAP4* fibroblasts. Co-staining for *COL3A1* and *CXCL14* was used to identify *COL3A1*/*CXCL14* fibroblasts, as this was predicted to be more sensitive than staining for either marker alone. Staining for *ACTA2* alone was used to identify *ACTA2* fibroblasts.

### Image acquisition and RNAscope signal quantification

RNAscope-stained sections were imaged by confocal microscopy using identical acquisition settings across samples and conditions. Anatomically distinct regions of reactive lymph nodes, including subcapsular sinus, T–B border, T-cell zone, perivascular regions and medullary sinus, were identified based on tissue architecture and marker distribution. CLL lymph nodes lacked germinal centers and therefore did not contain defined T cell zones or T-B borders. Acquired images were imported into ImageJ, where region-specific fields of view corresponding to the annotated anatomical compartments were manually cropped for quantitative analysis. For each RNAscope channel, fluorescence signal was thresholded uniformly across samples, and the area occupied by probe signal was measured. RNAscope signal was expressed as the percentage of total image area occupied by each probe channel within the cropped region. Identical thresholding and analysis parameters were applied across all samples and conditions. Quantified values were averaged per case, and each data point represents an independent lymph node sample.

### Spatial transcriptomics

#### Sample preparation and initial segmentation

FFPE lymph node sections were prepared by the Specialized Histopathology Core, Brigham and Women’s Hospital, for spatial transcriptomics on the 10x Genomics Xenium *In Situ* platform according to the manufacturer’s protocols. Probe hybridization, imaging, and initial data generation were performed by the Center for Cellular Profiling, Brigham and Women’s Hospital. Xenium data were generated using the Xenium Prime 5K human gene expression panel and processed using the Xenium Analyzer pipeline to produce per cell gene expression matrices and spatial coordinates. Cell segmentation and transcript assignment were performed using the Xenium Analyzer software following standard workflows. The three CLLs analyzed by spatial transcriptomics (cases CLL1, CLL3, and CLL8, Supplemental Table 1) contained 170,508, 270,761, and 349,696 cells, respectively. Median transcript counts were 510 transcripts per cell for CLL1, 513 for CLL2, and 286 for CLL3, indicating robust transcript detection and consistent data quality. Distributions of transcript counts per cell and additional dataset quality control metrics are provided in Supplementary Table 7.

#### Custom spatial analysis

Downstream spatial analyses were performed using custom scripts in R. Spatial distance analyses were performed using Xenium-derived cell centroid coordinates.

#### Spatial distance analysis of *MS4A1*(CD20)^+^ cells to CCL19 hotspots

*MS4A1*⁺*MYC⁺* cells were defined as cells with ≥3 *MS4A1* transcripts and ≥2 *MYC* transcripts. Here and in the analyses described below, *MS4A1⁺MYC⁻* cells were defined as cells with ≥3 *MS4A1* transcripts and no *MYC* transcripts. Proliferation center-associated *MS4A1*⁺*MYC⁺* cells were defined as the densest 40 percent of *MS4A1*⁺*MYC⁺* cells, based on the mean distance to their *k* nearest neighbors (*k* = 5). *CCL19* hotspots were defined as the top 1 percent of *CCL19* expressing cells within each sample. Euclidean distances were calculated from individual *MS4A1*⁺*MYC⁺* and *MS4A1⁺MYC⁻* cells to the nearest CCL19 hotspot. Statistical comparisons were performed within each CLL case using two-sided Wilcoxon rank-sum tests.

#### Spatial distance analysis of *MS4A1*(CD20)^+^*MYC⁺CCR7⁺* cells to CCL19 hotspots

*MS4A1^+^MYC⁺CCR7⁺* cells were defined as cells with cells with ≥3 *MS4A1* transcripts and any level of *MYC* and *CCR7* transcripts. *CCL19* hotspots were defined as the top 1 percent of *CCL19* expressing cells within each sample. Euclidean distances were calculated from individual *MS4A1^+^MYC⁺CCR7⁺* and *MS4A1^+^MYC^-^*cells to the nearest *CCL19* hotspot. Statistical comparisons were performed within each CLL case using two-sided Wilcoxon rank sum tests.

#### Spatial distance analysis of *MSA41(CD20)^+^MYC⁺ITGAX⁺ITGB2⁺* cells to *ICAM1⁺* macrophages

*MSA41^+^MYC⁺ITGAX⁺ITGB2⁺* cells were defined as cells with ≥3 *MS4A1* transcripts and any level of *MYC*, *ITGAX*, and *ITGB2* transcripts. To identify spatially clustered *MSA41^+^MYC⁺ITGAX⁺ITGB2⁺*, mean k nearest neighbor distances were calculated (k = 5) and the densest 40 percent of cells were designated as proliferation center-associated. *ICAM1⁺*macrophages were defined as cells with ≥2 CD68 transcripts and any level of *ICAM1* transcripts. Euclidean distances were calculated from individual *MSA41^+^MYC⁺ITGAX⁺ITGB2⁺* cells and *MS4A1^+^MYC^-^*cells to the nearest *ICAM1⁺*macrophage. Statistical comparisons were performed within each case using two-sided Wilcoxon rank sum tests.

#### Spatial distance analysis of *MS4A1CD20⁺MYC⁺TNFRSF13C(BAFFR)⁺* cells to *TNFSF13B(BAFF)⁺* macrophages

*MS4A1^+^MYC⁺TNFRSF13C⁺* cells were defined as cells with ≥3 *MS4A1* transcripts and any level of MYC and TNFRSF13C transcripts. To identify spatially clustered *MS4A1^+^MYC⁺TNFRSF13C⁺* cells, mean *k* nearest neighbor distances were calculated (*k* = 5) and the densest 40 percent of cells were designated as proliferation center-associated. *TNFSF13B*⁺ macrophages were defined as cells with ≥2 CD68 transcripts and any level of *TNFSF13B* transcripts. Euclidean distances were calculated from individual *MS4A1^+^MYC⁺TNFRSF13C⁺* and *MS4A1⁺MYC⁻* cells to the nearest *TNFSF13B⁺* macrophage. Statistical comparisons were performed within each case using two-sided Wilcoxon rank sum tests.

#### Radial spatial analysis of *LSGAL9* (GAL9) expression near *MS4A1(CD20)⁺MYC⁺*cells

Two methods were used. In **<u>method 1</u>** (Figure 6K), continuous radial profiling was performed to quantify the distribution of *LGALS9* expression around proliferation center–associated, clustered *MS4A1^+^MYC⁺*cells, using the same Xenium-derived cell centroid coordinates. Euclidean distances were calculated from each clustered cell to all surrounding cells within the tissue section. *LGALS9* expression was averaged within successive 20 µm radial distance bins extending from clustered *MS4A1^+^MYC⁺* cell and radial profiles were computed independently for each cell. Case-specific radial profiles were generated by averaging bin means across all clustered cells within each sample and visualized as a function of distance by connecting bin means without normalization. In **<u>method 2</u>** (inner versus outer shell analysis, Supplemental Figure 13C) spatial enrichment of *LGALS9* expression near proliferation centers was assessed using Xenium-derived cell centroid coordinates. *MS4A1^+^MYC⁺*proliferation center cells were defined as cells with ≥3 *MS4A1* and ≥2 *MYC* transcripts, and spatially clustered cells were identified as the densest 40 percent of *MS4A1^+^MYC⁺* cells based on mean k nearest neighbor distances (k = 5). For each clustered *MS4A1^+^MYC⁺*cell, Euclidean distances to all cells were calculated and *LGALS9* transcript signal was quantified within concentric radial distance shells. Inner and outer regions were defined as 10–200 µm and 200–500 µm from clustered *MS4A1^+^MYC⁺* cells, respectively. *LGALS9* values were normalized in each case to the mean signal in the inner (10–200 µm) shell. Statistical comparisons between inner and outer regions were performed in each case using paired Wilcoxon tests.

### Immunostaining

Immunostaining was performed on formalin-fixed paraffin-embedded sections with antibodies specific for MYC (Abcam, cat. #AB32072), Ki67 (Cell Signaling Technology [CST], cat. #9449T), CD20 (CST, cat. #48750T; or GeneTex, cat. #21310), COX-IV (CST, cat. #11967), CCR7 (R&D, cat. #MAB197), MIF (Sigma, cat. #HPA003868), CD74 (ThermoFisher, cat. #MA5-29128), Galectin9 (GAL9; CST, cat. #54330), CD44 (Abcam, cat. #ab189524). Antigen retrieval was performed using citrate (pH 6) or Tris-EDTA (pH 9) buffers (Vector Labs), as appropriate. Antibodies were diluted in signal stain antibody diluent (CST, cat. #8112L). Signal amplification was performed with horseradish peroxidase (HRP) based technology using Novolink polymer (cat. #RE7290-CE, Leica Biosystems), SignalStain Boost (CST, cat. #8125P), and anti-goat IgG-HRP (Vector labs, cat. PI-9500). For HRP-based fluorescent dye labeling, we used Alexa Fluor 488 tyramide (Thermo Fisher, cat. #B40953) or iFluor styramide 555 or 594 (AAT Bioquest, cat. #45027 and 45035). In Figure 4E, to simplify the staining process, the position of CLL proliferation centers was inferred by immunostaining for MYC in a 4 μM serial section prepared from the same CLL case.

### Microscopy imaging

A Zeiss LD LCI Plan Apo 25x/0.8 multi-immersion DIC II objective lens was used for single point scanning confocal imaging on a Zeiss Axio Observer Z1 with an LSM980 scan head. Imaging was performed in an environmental enclosure using a galvanometer (galvo) scanner. Fluorophores covered the range of 405-840nm. All images were processed using Zen Blue Acquisition Software.

For immunofluorescence microscopy imaging to validate ligand-receptor pairs, we used a Leica Stellaris 8 Falcon confocal microscope with a Leica Plan Apo 20x/0.75 air objective lens. Images were acquired using a resonant scanner with a pixel dwell time of 1.58μs 2x line averaging, and a pixel size of 189.39nm x 189.39nm (set as optimal for confocal detection) by Leica LAS X Acquisition Software and exported as LIF files. For all images, the pinhole diameter was 1AU (set as optimal for confocal detection). Tile scans were acquired across the desired areas, and the corresponding images were merged automatically in the LAS X software. Fluorophores were excited using a white light laser (WLL) with AOTF modulation that was tunable from 440-790nm. Emissions were collected using HyS and HyDX spectral detectors with collection windows optimized per fluorophore from 405-840nm.

### Microscopic image analysis

#### Measurement of MYC/Ki67 colocalization in proliferation centers

Proliferation centers were defined as clusters of at least five neighboring CD20⁺MYC⁺ cells. Within each proliferation center, Ki67⁺ cells were quantified and the proportion of CD20⁺MYC⁺ cells co-expressing Ki67 was determined.

#### Measurement of combined *in situ* hybridization and immunostain signal strength

To quantify *in situ* hybridization signal intensities in CLL cells within and outside of proliferation centers, we outlined the area encompassing proliferation centers (defined by the presence of clusters of MYC^+^ cells) with the ImageJ polygon area tool. Signals were then quantified independently for MYC (detected by immunofluorescent staining with anti-MYC antibody) and the fibroblast-specific RNAscope probes *CXCL14*/*COL3A1*, *CCL21*, *CCL19*, and *ACTA2* using CellProfilerTM (version 4.2.1). To compare marker gene expression inside and outside of proliferation centers, we measured the average signals of 4 randomly selected areas external to proliferation centers and quantified *in situ* hybridization signals as above.

#### Imaging of ligand-receptor pairs

To study colocalization of ligand-receptor pairs, we performed multiplex fluorescence confocal microscopy on CLL proliferation centers. Two ligand-receptor pairs, MIF-CD74 and GAL9-CD44, were stained in sections prepared from FFPE LN samples and tested for colocalization using a pipeline in ImageJ Fiji (version 1.53q). Briefly, we performed background subtraction and manual thresholding of individual signals for each channel using rolling ball radius followed by conversion to binary images. The same thresholds were applied to each channel in MYC-high and MYC-low areas of the same stained tissue section. After merging images of different channels, locations where signals of 2 channels coincided were considered areas of colocalization. The percentage of colocalization was defined as the percentage of the area of ligand staining that overlapped with the area of staining for the corresponding receptor.

For studies involving MIF-CD74, because cancer cells are expected to be “receivers”, we assessed colocalization of MIF (ligand) and CD74 (receptor) on the surface of MYC^+^CD20^+^ CLL cells. MYC^+^CD20^+^ cells were defined by expanding the area of the MYC signal using the ImageJ dilate function and selecting cells with overlapping MYC and CD20 signals. We then compared ligand and receptor colocalization in CD20+ cells that were outside of proliferation centers. Because GAL9 is secreted and predicted to act on CD44 expressed on immune cells, we measured colocalization GAL9 and CD44 on cells located within and outside of circle with a radius of 30 μm centered on MYC^+^CD20^+^ cells. We selected 30 μm as a cutoff because secreted proteins (i.e. cytokines) have been reported to robustly activate cognate receptors on neighboring cells up to 30 μm away (see Thurley, K. et al., ref. #59).

### Spheroid culture and proliferation assays

Primary peripheral blood CLL cells derived from IGHV-mutated (n = 3) or IGHV-unmutated (n = 3) cases were thawed and immediately diluted dropwise into cold B-cell medium (ExCellerate, xeno-free) supplemented with 20% fetal bovine serum (FBS) and 1% penicillin/streptomycin, followed by incubation for 20 min in the dark. Cells were then centrifuged for 5 minutes at 500 × g, the supernatant was discarded, and pellets were resuspended in PBS. Cells were counted using Cellometer^TM^ cell counting chambers. Subsequently cells were labeled with 4 µM CellTrace Violet (CTV; Thermo Fisher Scientific; cat. # C34571) according to the manufacturer’s instructions. Labeled cells were centrifuged at 200 × g for 5 min and pellets resuspended in B-cell medium (ExCellerate, xeno-free) supplemented with 2% FBS and 1% penicillin/streptomycin, and seeded at 100,000 cells per well in 300 µL of medium in ultra-low attachment round-bottom 96-well spheroid plates (Corning, 4515) to promote three-dimensional spheroid formation. Cells were cultured for 7 days either in the absence of added stimuli (unstimulated) or in the presence of the indicated growth factors. Where indicated, growth factors were added at the following final concentrations: IL-2 (25 ng/mL; PeproTech; cat. #200-02), IL-15 (25 ng/mL; PeproTech; cat. #200-15), IL-21 (25 ng/mL; PeproTech; cat. # 200-21), IL-4 (30 ng/mL; PeproTech; cat. # 200-04), MEGA-CD40L (50 ng/mL; Enzo Life Sciences; NC9975949), and CpG ODN 2006 (1 µg/mL; InvivoGen; cat. #NC9205905). Growth factors were added at the start of the culture and maintained throughout the 7-day assay period. For CD74 functional blocking experiments, a mouse anti-human CD74 blocking antibody (Clone LN2, SouthernBiotech; cat. #9775-01) was added at a final concentration of 1 µg/mL at the start of the assay and maintained throughout the culture period. The corresponding mouse IgG1 κ isotype control antibody (BioLegend; cat. # 400102) was used at the same concentration.

After 7 days of culture, CLL cells were harvested from spheroid cultures and analyzed by flow cytometry. Cells were first incubated with Human TruStain FcX Fc receptor blocking solution (BioLegend; cat. # 422302) according to the manufacturer’s instructions. Cells were stained with APC anti-human CD5 antibody (BioLegend; cat. #364016) and PE anti-human CD19 antibody (BioLegend; cat. #392506) to identify CLL cells. Cell viability was evaluated using Zombie NIR Fixable Viability Dye (BioLegend; cat. #77184) and apoptosis was assessed using Annexin V PE-Cy7 (BioLegend; cat. #640951) in Annexin V binding buffer according to the manufacturer’s instructions. Cell proliferation was assessed based on CellTrace Violet dilution within gated live (Zombie NIR⁻) Annexin V⁻CD5⁺CD19⁺ CLL cells. Data were acquired on a BD FACSymphony A1 flow cytometer (BD Biosciences) using standard compensation controls and analyzed using FlowJo software. The percentage of divided cells was determined based on loss of CTV signal relative to the undivided population. Each data point represents an independent CLL case. Spheroid formation and growth were monitored by brightfield microscopy and spheroid area was quantified using ImageJ.

### Statistical analysis

Statistical analysis was conducted using Prism 9 (GraphPad Software). Prior to analysis, data were evaluated for a Gaussian distribution using either the Kolmogorov-Smirnov test with the Dallal-Wilkinson-Lillie correction for P value or the Shapiro-Wilk normality test. If data were normally distributed, the Student’s t-test for two groups or one-way ANOVA and Dunnett’s test for two or more groups were used as appropriate. If the data were not normally distributed, the Mann-Whitney test was used to determine the significance of differences between two groups, the two-sided Wilcoxon signed-rank test was used for paired comparisons, and Kruskal-Wallis and Dunn’s tests were used for multiple comparisons for two or more groups as appropriate. Data are presented as mean ± SEM. Statistical significance was defined as *P* <0.05 and is indicated as ns (non-significant), *P <0.05, **P <0.01, or ***P <0.001. Sample size was not predetermined through statistical methods.

### Data Sharing Statement

scRNA-seq data sets are available through GEO: GSE261917.

