## Supplemental Figures for "CCL19⁺ fibroblasts define a proliferative niche in chronic lymphocytic leukemia"

### Supplemental information

#### Table legends

**Supplementary Table 1:** Pathologic characteristics of lymph nodes selected for analysis.

**Supplementary Table 2:** Cell type defining markers, and sequencing and quality control metrics.

**Supplementary Table 3:** Differential gene expression in reactive and CLL lymph node cells. Related to Figure 1C.

**Supplementary Table 4:** Differential gene expression in reactive and CLL lymph node $CD45^{+}$ cells. Related to Figure 2B, and Supplemental Figure 2B-C.

**Supplementary Table 5:** Differential gene expression of $CD45^{-}$ cells, fibroblast subsets, and blood and lymphatic endothelium in reactive and CLL lymph nodes. Related to Figure 4B and D, and Supplemental Figures 6E, 8C, and 8H.

**Supplementary Table 6:** CellPhoneDB-predicted ligand-receptor interactions between $MYC^{+}$ and $MKI67^{+}$ proliferating CLL cells and CCL19-high/CCL21-low fibroblasts, together with associated differential expression patterns. Related to Figure 5F and G and Supplemental Figure 9C

**Supplementary Table 7:** CellChat, CellPhoneDB and NicheNet-predicted ligand-receptor interactions between $MYC^{+}$ and $MKI67^{+}$ proliferating CLL cells and $CD45^{+}$ immune cells, annotation of genes present in the Xenium Prime Human 5K panel, and Xenium metrics. Related to Figure 6B-G, and Supplemental Figure 12.

#### Supplemental Figures

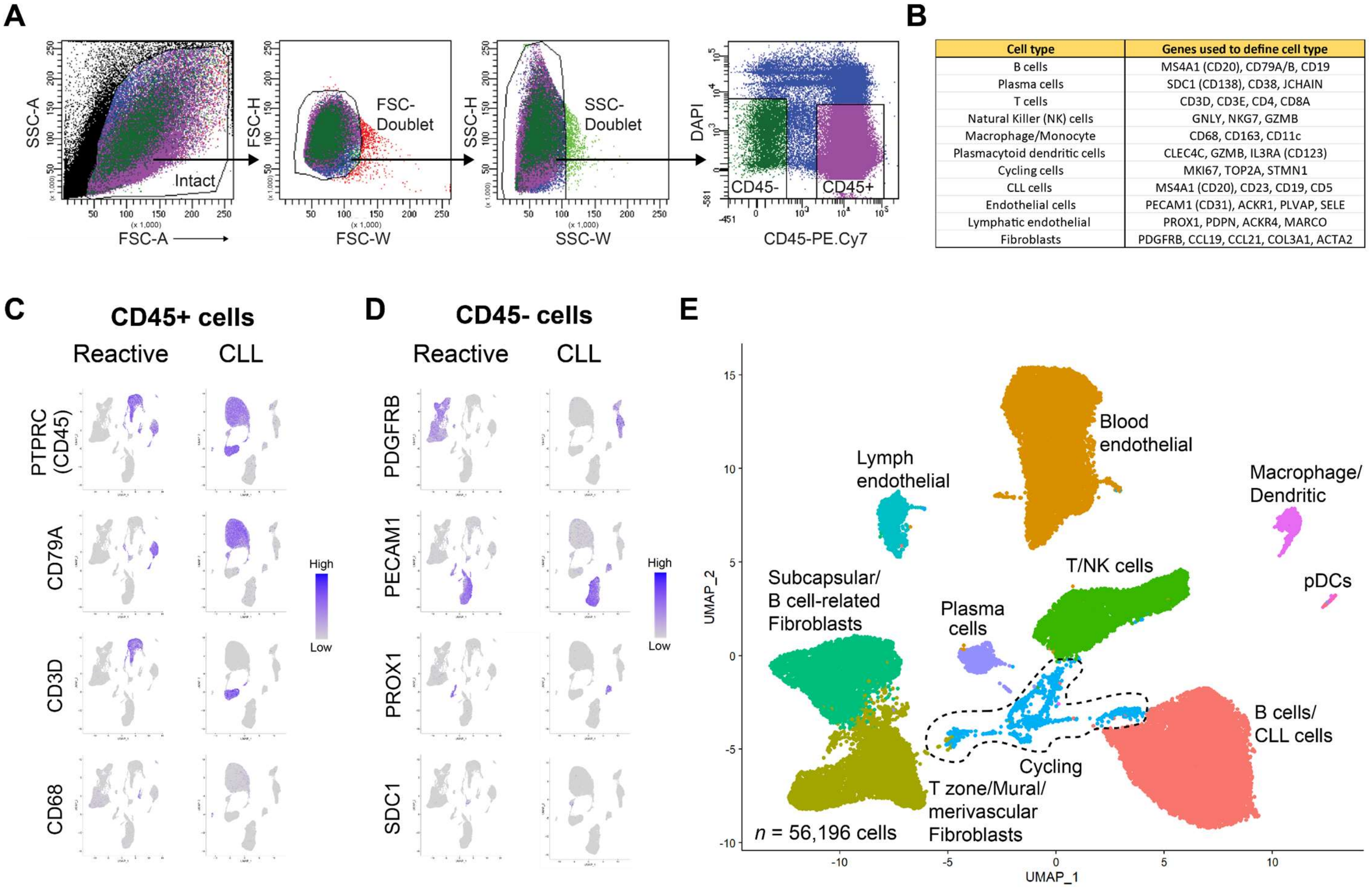


| Cell type | Genes used to define cell type |
|---|---|
| B cells | MS4A1 (CD20), CD79A/B, CD19 |
| Plasma cells | SDC1 (CD138), CD38, JCHAIN |
| T cells | CD3D, CD3E, CD4, CD8A |
| Natural Killer (NK) cells | GNLY, NKG7, GZMB |
| Macrophage/Monocyte | CD68, CD163, CD11c |
| Plasmacytoid dendritic cells | CLEC4C, GZMB, IL3RA (CD123) |
| Cycling cells | MKI67, TOP2A, STMN1 |
| CLL cells | MS4A1 (CD20), CD23, CD19, CD5 |
| Endothelial cells | PECAM1 (CD31), ACKR1, PLVAP, SELE |
| Lymphatic endothelial | PROX1, PDPN, ACKR4, MARCO |
| Fibroblasts | PDGFRB, CCL19, CCL21, COL3A1, ACTA2 |

**Supplemental Figure 1. Identification of main cell types in reactive and CLL LNs by scRNA-seq.**
**A)** Flow cytometry gating strategy used to sort CD45$^+$ or CD45$^-$ cells. **B)** Table showing genes used to define main cell lineages. **C, D)** Expression of lineage-specific genes mapped onto scRNA-seq UMAPs of integrated CD45$^+$(C) and CD45$^-$ (D) cells from reactive and CLL LNs. **E)** UMAP plot of integrated scRNA-seq data sets obtained from CD45$^+$ and CD45$^-$ cells from reactive (N=3) and CLL (N=3) LNs.

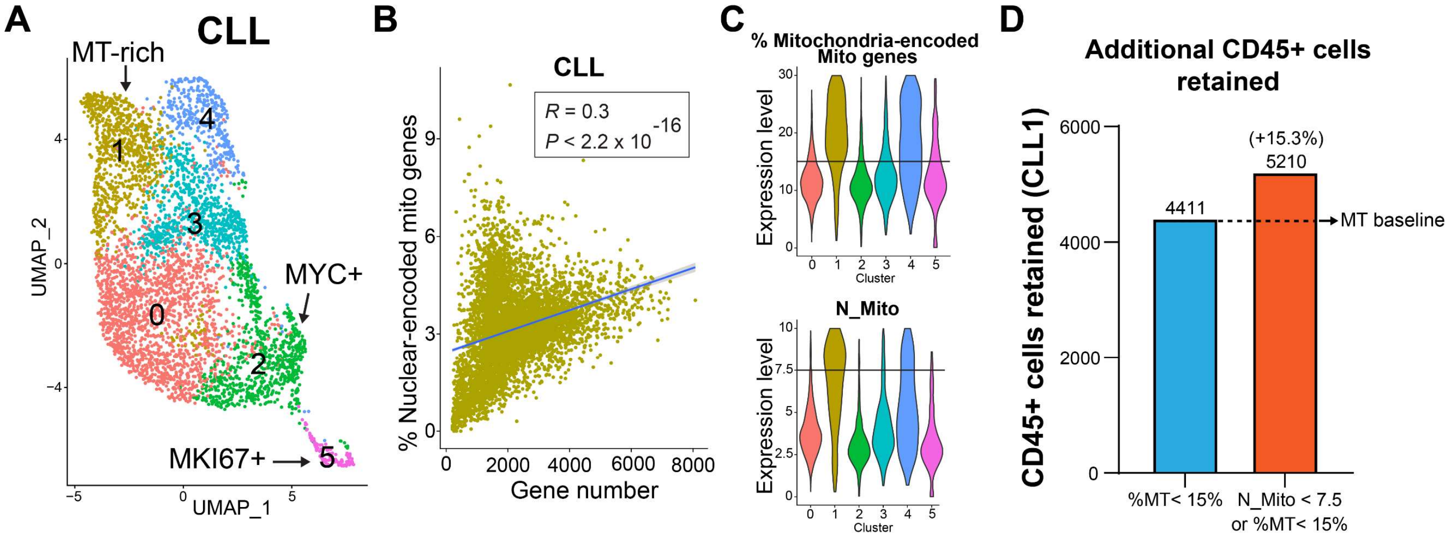


**Supplemental Figure 2. Rationale for retention of proliferating, mitochondrion-rich CLL cells using an alternative scRNA-seq filtering method, N_Mito.**

**A)** UMAP plot of cells from CLL case 1 showing *MKI67*$^{+}$ (cluster 5) and *MYC*$^{+}$ (cluster 2) clusters and a distinct cluster defined by high levels of mitochondrial gene-encoded transcripts (cluster 1, designated "MT-rich"). Mitochondrial filtering was not applied to this UMAP. **B)** Correlation between total gene transcripts and nucleus-encoded mitochondrial gene transcripts detected in CLL cells from A). **C)** Violin plot showing % expression of MT genes (Top) or N_Mito score (bottom) in each cluster shown in A. Horizontal lines represent cutoff thresholds (N_Mito < 7.5 or MT < 15%) used to filter cells. Note that the N_Mito cutoff retains more cells from clusters 2 and 5 (*MYC*$^{+}$ and *MKI67*$^{+}$, respectively). See Supplemental Methods for additional details. **D)** Bar plot showing the number of CD45$^{+}$ cells retained in CLL case 1 after standard %MT filtering alone (%MT < 15%) or after retaining cells that passed either mitochondrial gene filter (N_Mito < 7.5 or %MT < 15%).

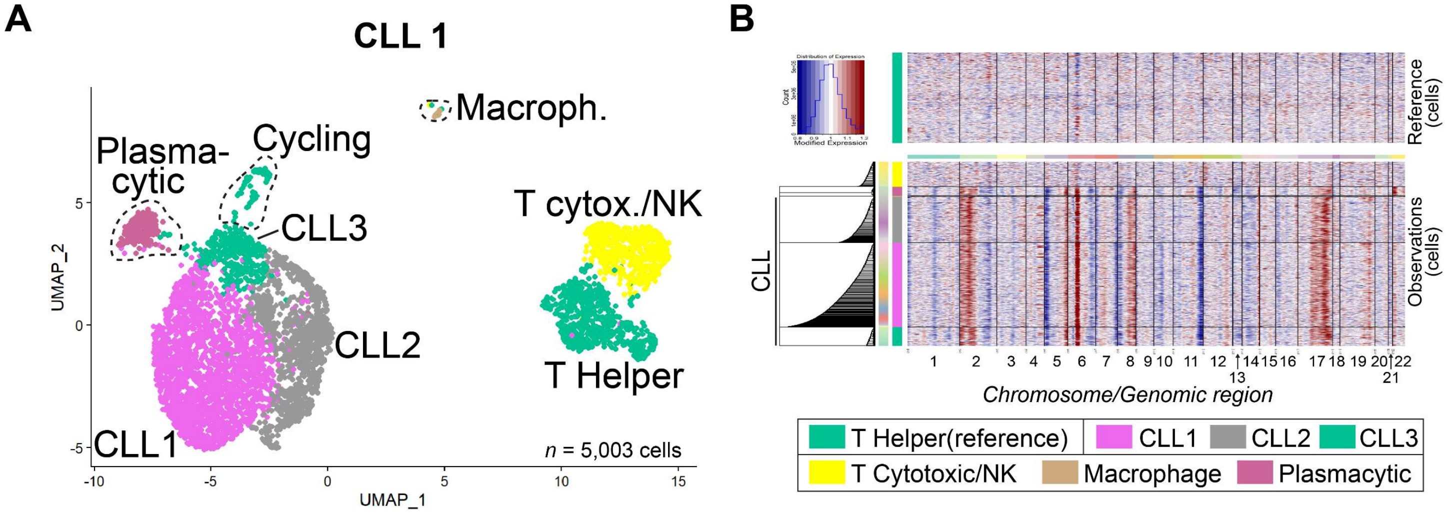


**Supplemental Figure 3. InferCNV analysis of CLL1 LN cells**.

**A)** UMAP of all CD45$^+$ cells from the CLL1 lymph node immune cells showing malignant B-cell subclusters, including cycling and plasmacytic populations. **B)** InferCNV analysis of the same CD45$^+$ CLL1 lymph node cells. Note that plasmacytic cells and CLL cells share the same inferred copy number variations (CNVs), whereas non-malignant immune cells do not.

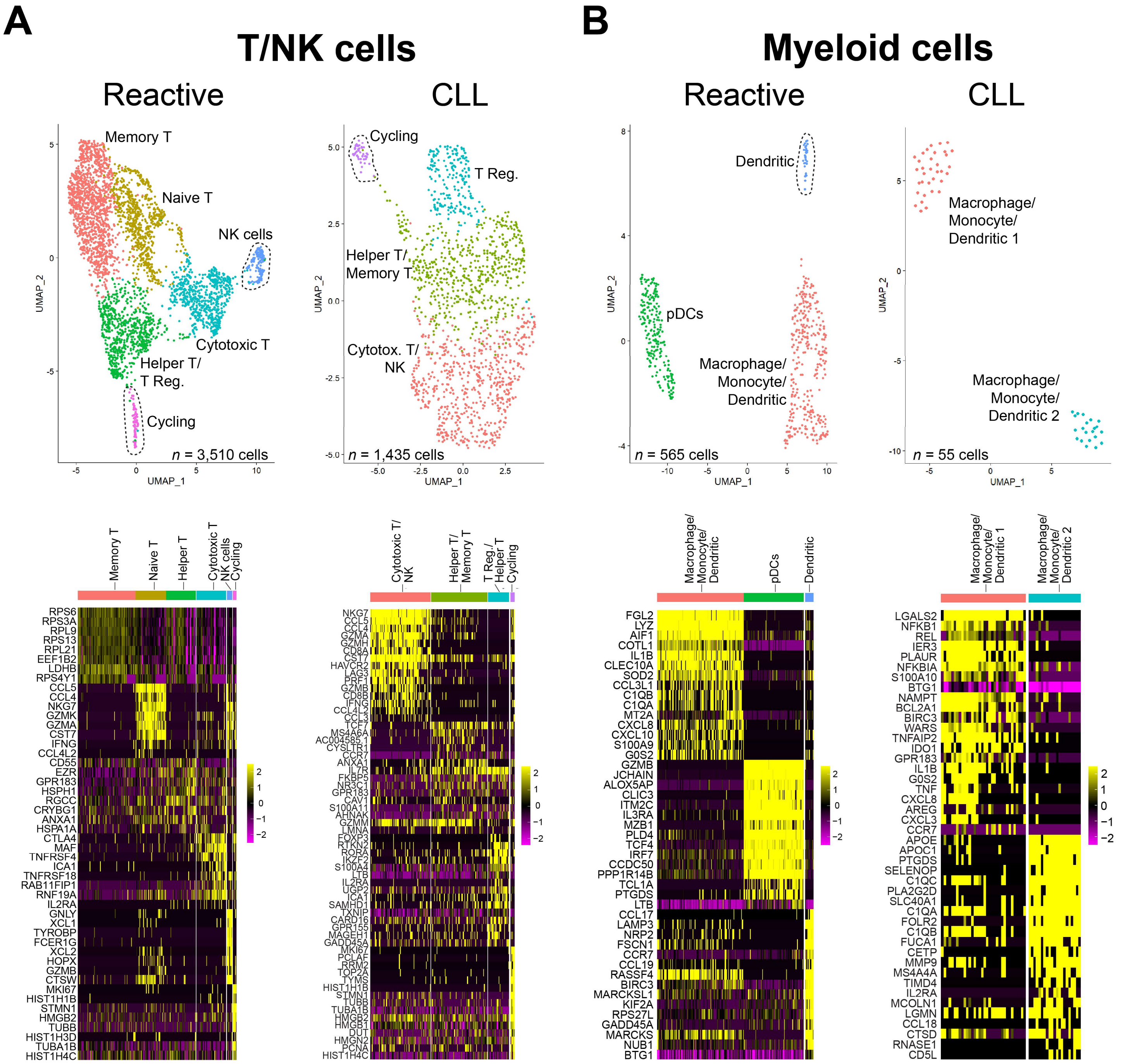


**Supplemental Figure 4. T/NK and myeloid cell subsets in reactive and CLL LNs**.

**A)** *Top*: UMAP plots showing clusters identified through integrated analyses of T/NK lineage cells from reactive and CLL lymph nodes. *Bottom*: heatmaps displaying top differentially expressed genes defining the clusters shown above. **(B)** Top: UMAP plots showing clusters identified through integrated analyses of myeloid lineage cells from reactive and CLL lymph nodes. Bottom: heatmaps displaying top differentially expressed genes defining the clusters shown above.

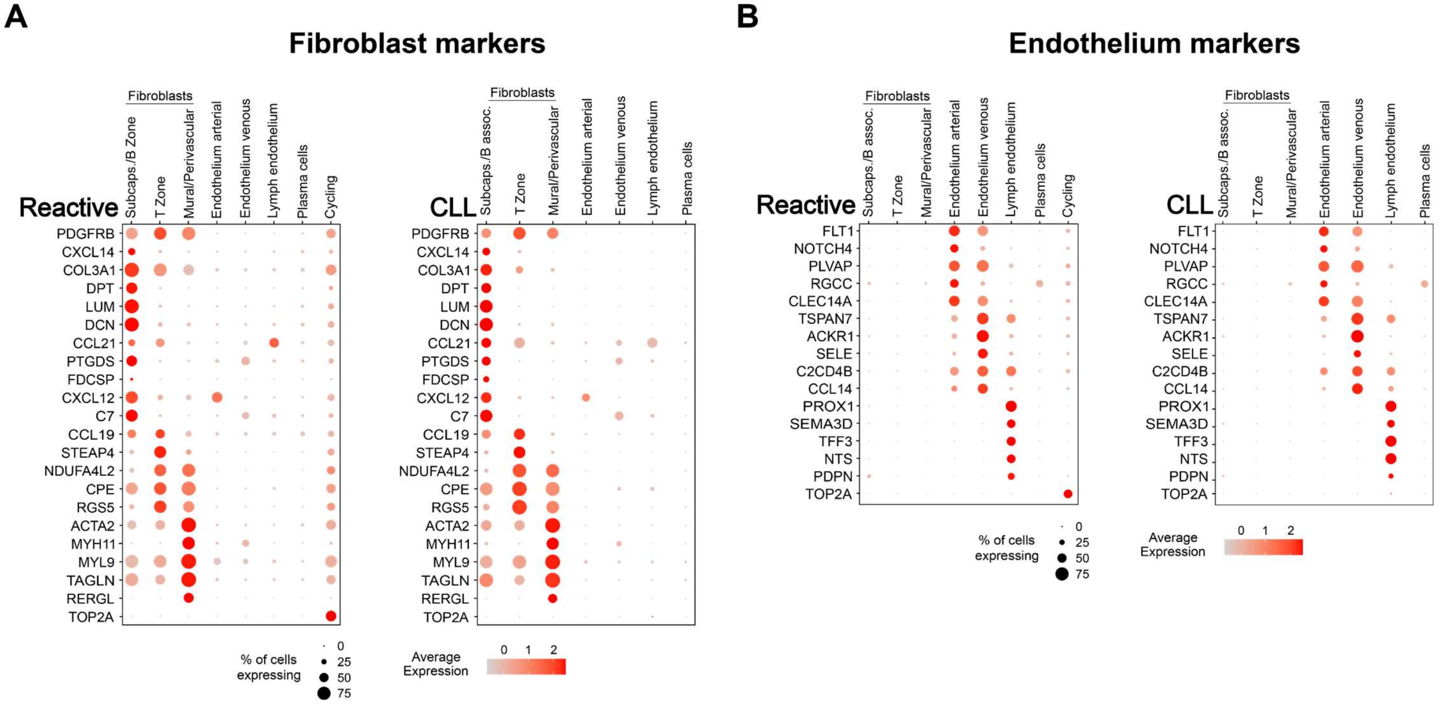


**Supplemental Figure 5. Differentially expressed genes in LN fibroblasts and endothelium.**
Dot plots displaying expression of fibroblast **(A)** and blood and lymph endothelium **(B)** cluster-specific genes. Clusters correspond to those identified in the UMAP plots shown in Figure 4A.

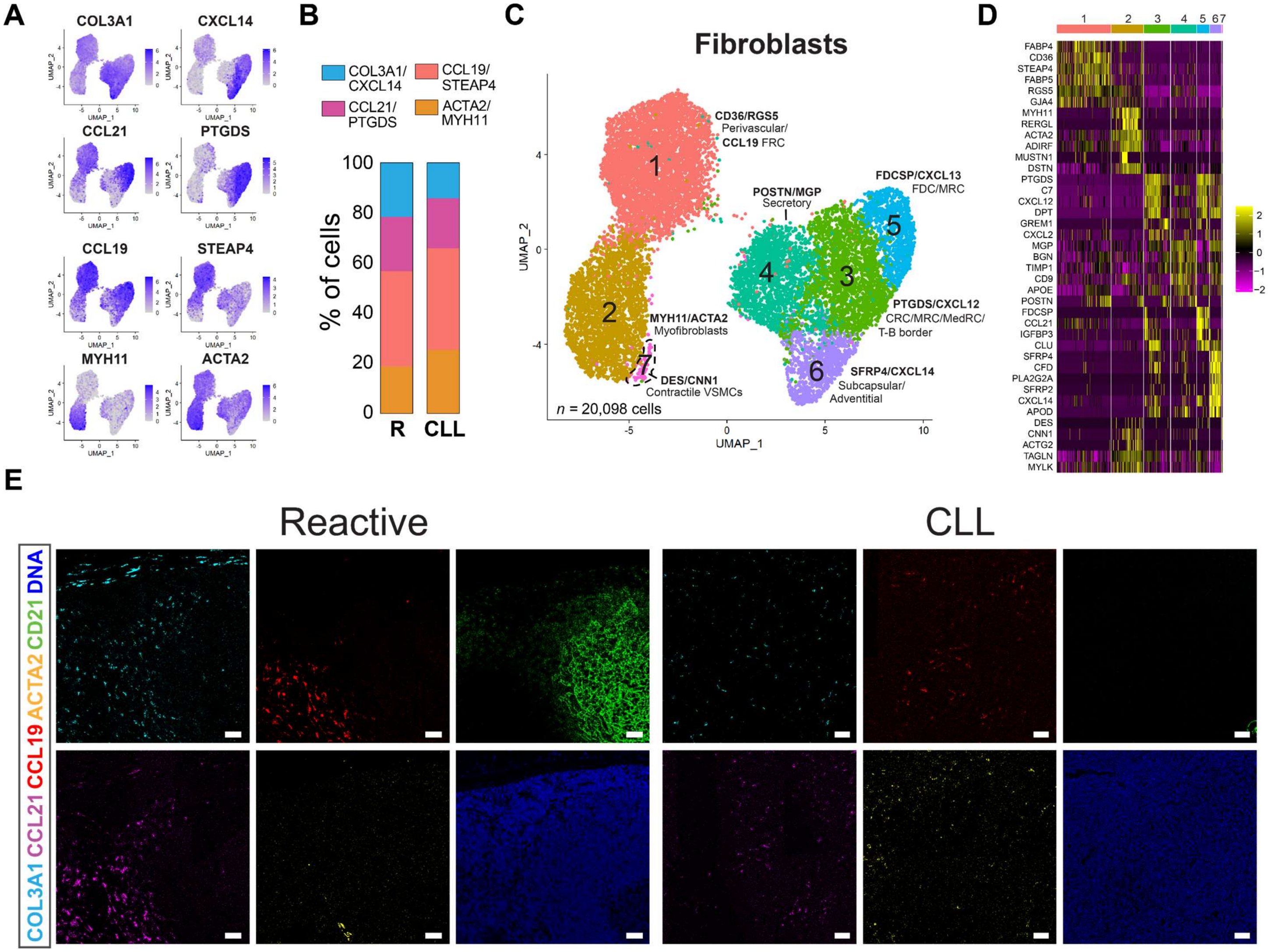


**Supplemental Figure 6. Major fibroblast types and subtypes in reactive CLL LNs**.

**A)** UMAP projections of *COL3A1*, *CXCL14*, *CCL21*, *PTGDS, CCL19*, *STEAP4* and *ACTA2*, *MYH11*, expression in integrated fibroblasts from reactive and CLL LNs. **B)** Percentage of the four main fibroblast types in reactive and CLL nodes based on integrated analysis. **C)** UMAP plots displaying the integrated scRNA-seq datasets of all fibroblasts from reactive and CLL LNs at a higher resolution, displaying 7 specific fibroblast subtypes. **D)** Heatmaps showing top differentially expressed genes in the 7 main fibroblast subtypes of panel D. **E)** Immunofluorescence confocal images showing individual channels derived from insets of Figure 4E. Scale bars, 50μm.

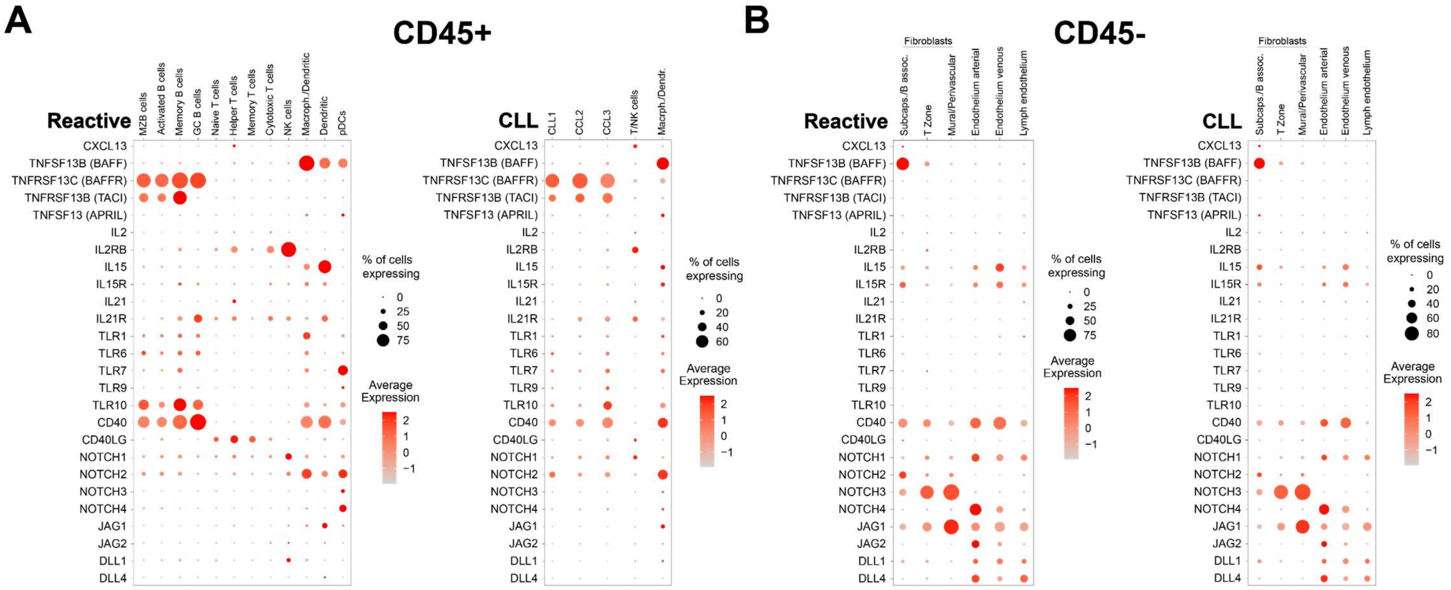


**Supplementary Figure 7. Expression of genes previously linked to CLL pathobiology in CD45⁺ and CD45⁻ cells isolated from reactive and CLL LNs.**

**A, B)** Dot plots displaying expression of genes linked to CLL pathobiology in CD45⁺ cells (A; see also Figure 2A) and CD45⁻ cells (B; see also Figure 4A).

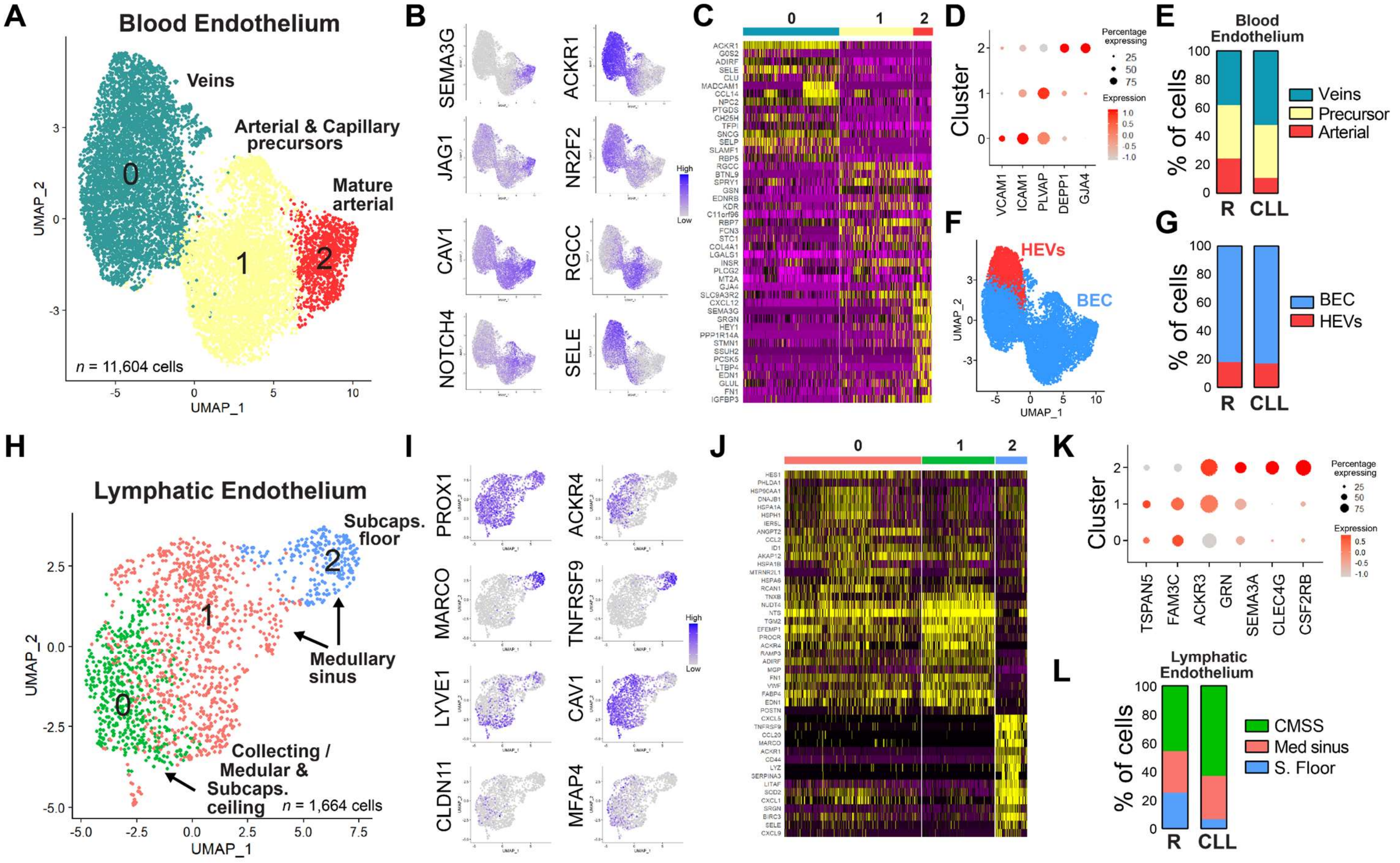


**Supplementary Figure 8. Blood and lymphatic endothelial cell populations in reactive and CLL LNs.**
**A)** UMAP plot displaying blood endothelial cell types identified in integrated analysis of reactive and CLL LNs. **B)** UMAP plots showing superimposed expression of genes used define the endothelium clusters in panel A. **C)** Heatmaps showing top differentially expressed genes in main blood endothelial cell clusters identified in panel A. **D)** Dot plots showing expression of genes that define specific blood endothelial cell clusters identified in panel A. **E)** Percentage of main blood endothelial cell types identified in reactive and CLL LNs in panel A). **F)** UMAP plot showing high endothelial venule endothelial cells (red) and other blood endothelial cells (blue) in the UMAP plot from panel A. **G)** Percentage of high endothelial venule endothelial cells and blood endothelial cells in reactive and CLL LNs. **H)** UMAP plot displaying main lymphatic endothelial cell types in integrated analysis of reactive and CLL LNs. **I)** UMAP plots showing superimposed expression of genes used to define the lymphatic endothelial cell clusters in panel F. **J)** Heatmaps showing top differentially expressed genes in lymphatic endothelial cell clusters identified in panel F. **K)** Dot plots showing expression of genes that define specific lymphatic endothelial cell clusters identified in panel F. **L)** Percentage of lymphatic endothelial cell cluster types identified in reactive and CLL LNs in panel F. BEC, blood endothelial cells; HEV, high endothelial vein endothelial cells; CMSS, collecting, medullary, and subcapsular ceiling lymphendothelial cells; Med sinus, medullary sinus lymphendothelial cells; S. Floor, subcapsular floor lymphendothelial cells.

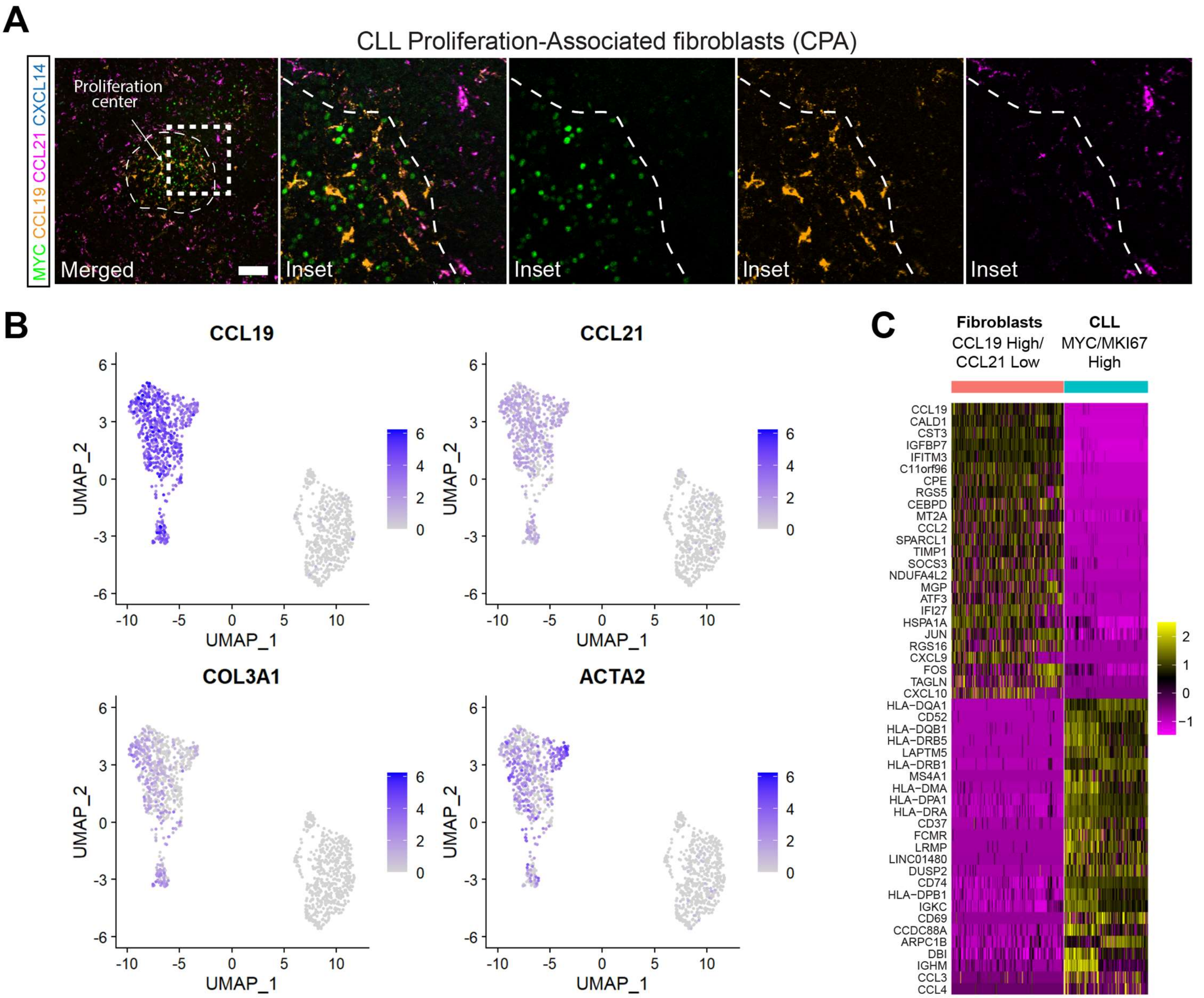


**Supplementary Figure 9. CPA fibroblast identification, fibroblast interactions with proliferating CLL cells, and differentially expressed genes.**

**A)** Confocal immunofluorescence images of MYC protein and *CCL19*, *CCL21*, and *COL3A1*/*CXCL14* transcripts from Figure 5A showing an inset with separate channels. Note that *CCL21* expression decreases and *CCL19* expression increases from the border zone to the interior of the PC. Scale bar, 100μm. **B)** CCL19-High/CCL21-Low fibroblasts (putative CPAs) interaction with MYC/MKI67-High CLL cells from the 3 integrated CLL LNs. **C)** Top differentially expressed genes in CCL19-high/CCL21-low fibroblasts and MYC/MKI67-High CLL cells from Figure 5E.

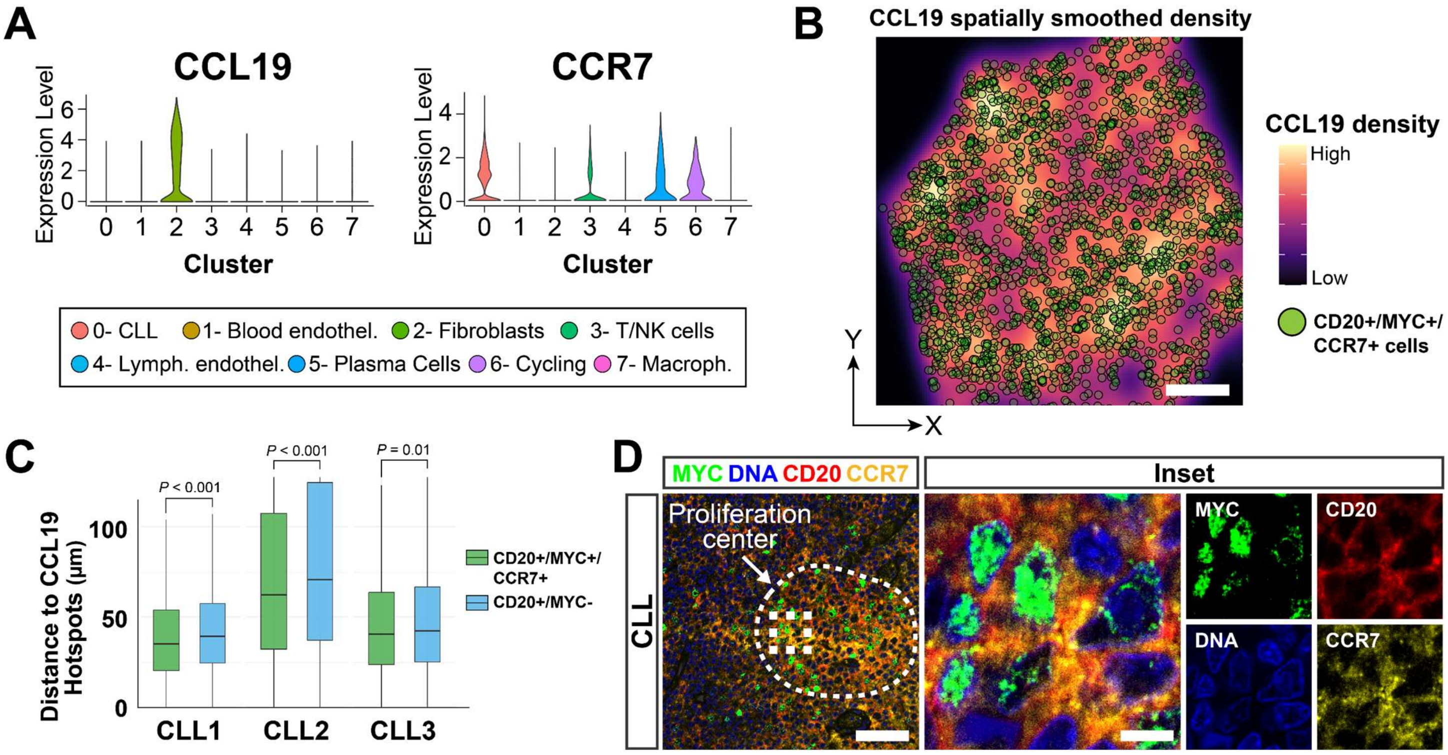


**Supplementary Figure 10.** ***CCR7-CCL19*** **interaction between CLL cells and** ***CCL19*****-High fibroblasts.**
**A)** Violin plots showing normalized expression levels of *CCL19* and *CCR7* across major cell populations from three integrated CLL LN scRNA-seq datasets. **B)** Spatial transcriptomics showing smoothed spatial density maps of *CCL19* expression overlaid with *MSA41*$^+$*MYC*$^+$*CCR7*$^+$ cells (green). Cells were slightly enlarged for visualization purposes. Scale bar, 400 μm. **C)** Average distances from *MSA41*$^+$*MYC*$^+$*CCR7*$^+$ cells and *MSA41*$^+$*MYC*$^-$ cells to CCL19-high “hotspots” (top 1% of signal) measured across 3 CLL LNs. Statistical comparisons were performed within each CLL case using two-sided Wilcoxon rank sum tests. **D)** Confocal immunofluorescence microscopy of a CLL proliferation center showing colocalization of CD20$^+$MYC$^+$ CLL cells, and CCR7. Scale bars: 100 μm (merged image), 5 μm (inset).

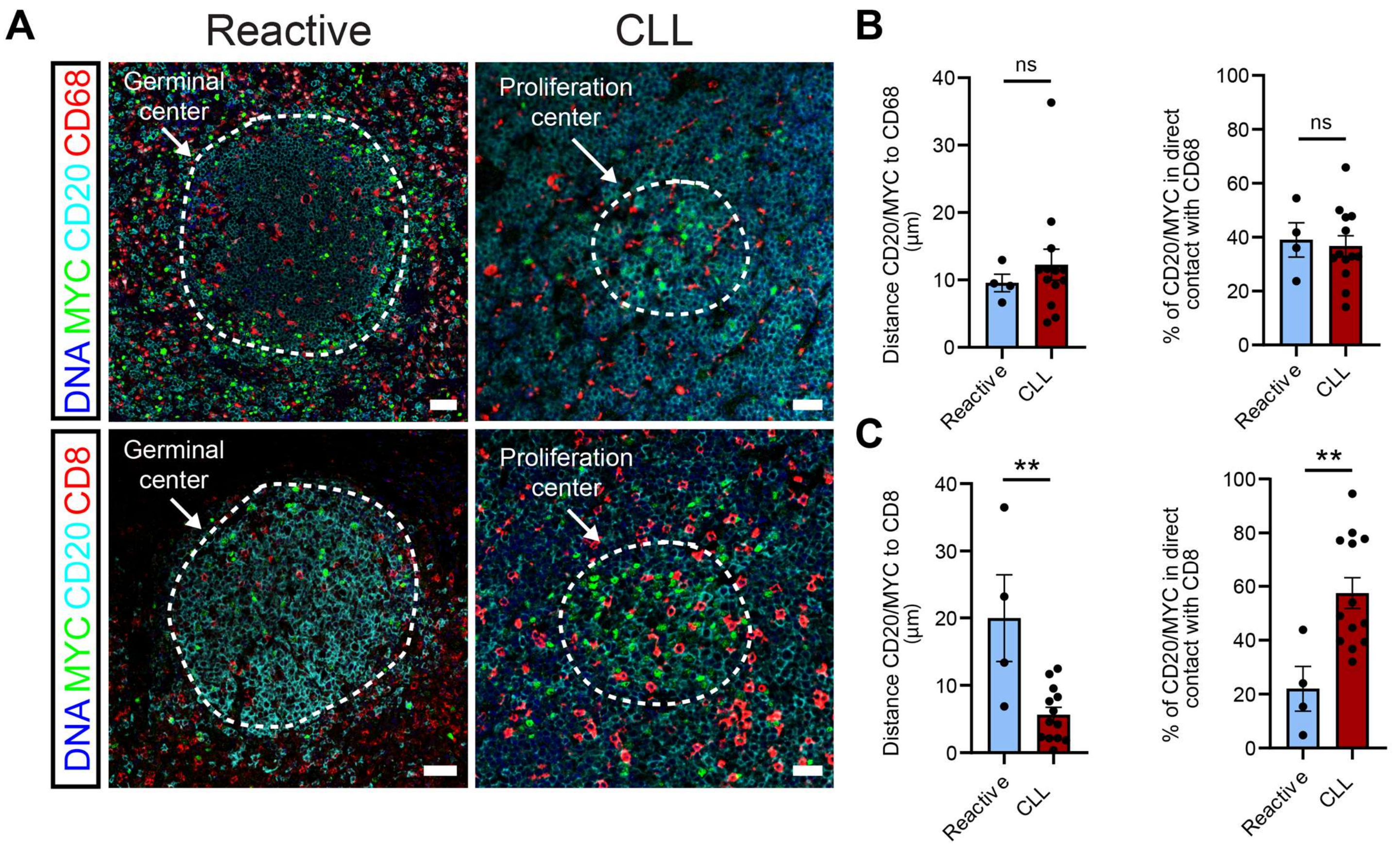


**Supplemental Figure 11. Distances between CD20+/MYC+ cells and macrophages and T cytotoxic cells in CLL and reactive LNs.**

**A)** Multiplex immunofluorescence microscopy depicting CD20, MYC and CD68 (macrophage marker, top) or CD8 (cytotoxic T cell marker, bottom) within CLL proliferation centers and reactive germinal centers. Scale bars, top, 50 µm and 30 µm, and bottom, 50 µm and 25 µm, for reactive and CLL, respectively. **B)** Distance between CD20+/MYC+ cells and CD68+ cells in CLL proliferation centers and reactive germinal centers (left), and percentage of CD20+/MYC+ cells in direct contact with CD68+ cells (right). Each dot represents the average distance of at least 50 cells from each LN. **C)** Same as B) but distances and percentage of total cells in direct contact are compared to CD8 cytotoxic T cells.

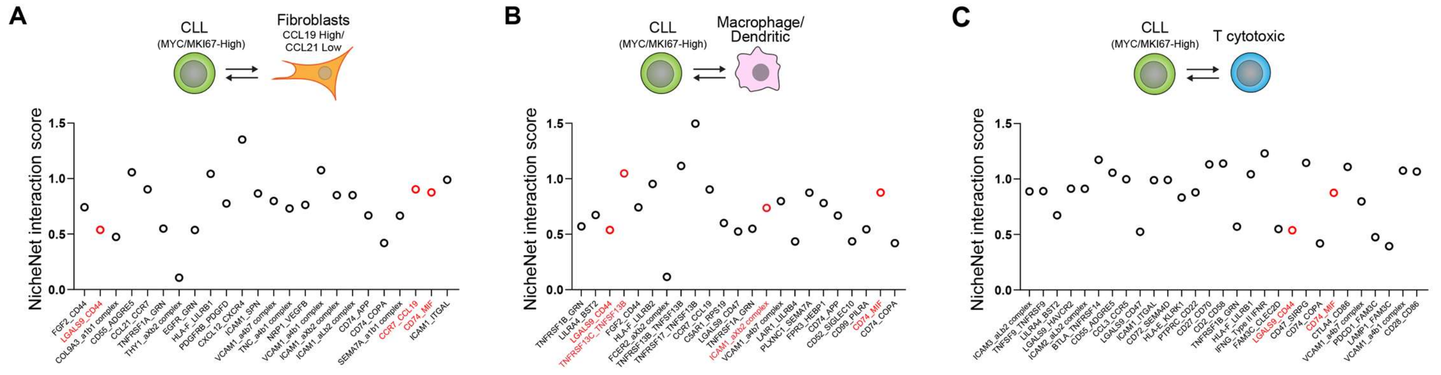


**Supplemental Figure 12. NicheNet ligand-receptor interaction scores between MYC/MKI67-High CLL cells cells and CCL19-High/CCL21-Low fibroblasts, macrophages, and cytotoxic T cells.**

**A-C)** NicheNet interaction score of top 20 CellPhoneDB-predicted ligand-receptor pairs expressed by proliferating MYC/MKI67-High CLL cells and (A) *CCL19*-high/*CCL21*-low fibroblasts (putative CPAs), (B) macrophages, and (C) cytotoxic T cells. Ligand-receptor pairs highlighted in red correspond to interactions that were further examined by in situ staining, spatial transcriptomics, and/or growth assays.

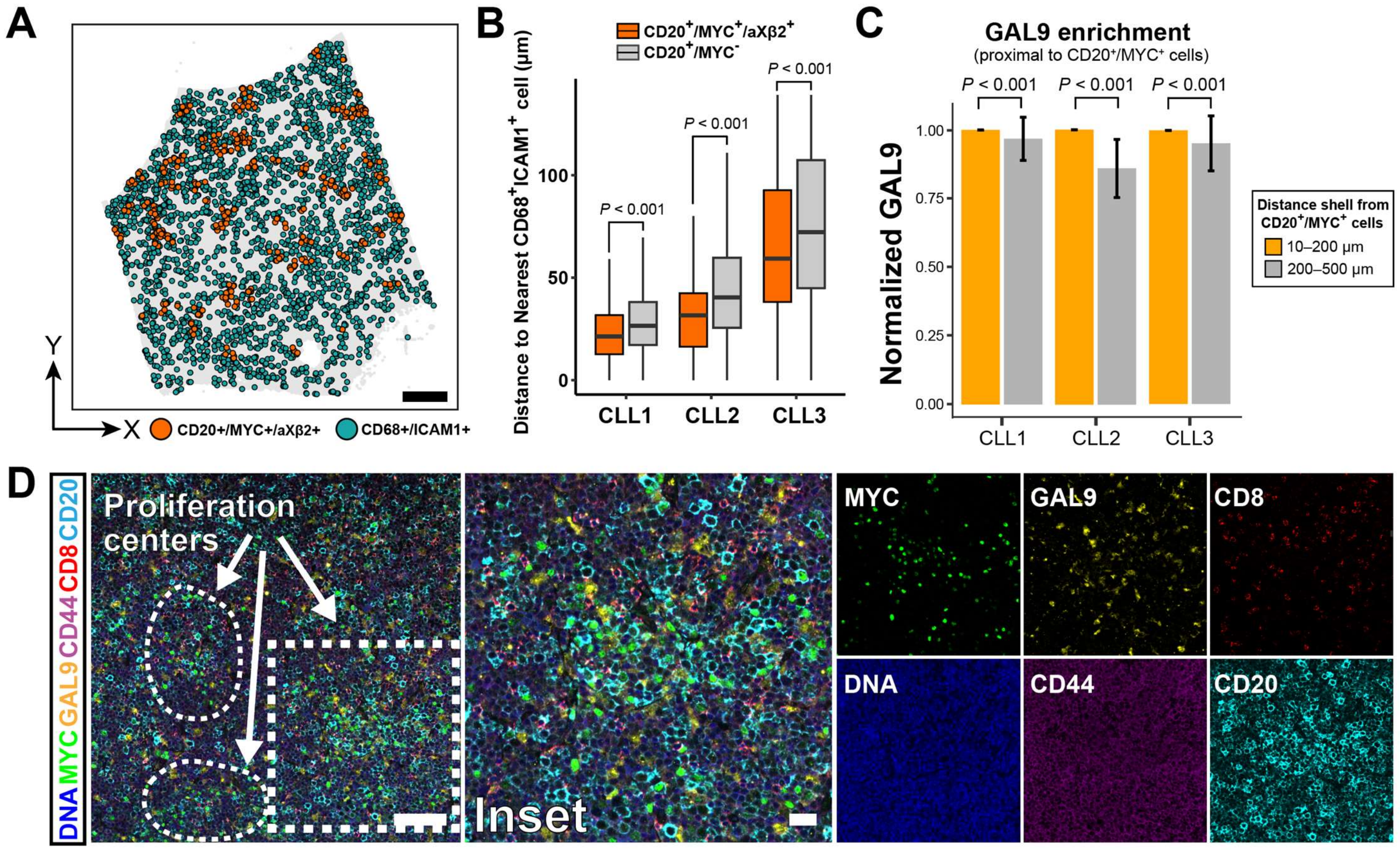


**Supplemental Figure 13. Spatial association of ICAM1$^{+}$CD68$^{+}$ macrophages and GAL9 enrichment within CLL proliferation centers.**

**A)** Spatial distribution of *MS4A1*$^{+}$*MYC*$^{+}$*ITGAX*$^{+}$*ITGB2*$^{+}$ CLL cells and *ICAM1*$^{+}$*CD68*$^{+}$ macrophages in a CLL LN. Cells were slightly enlarged for visualization purposes. Scale, 400 μm. **B)** Nearest-neighbor analysis comparing distances between *MS4A1*$^{+}$*MYC*$^{+}$*ITGAX*$^{+}$*ITGB2*$^{+}$ and *MS4A1*$^{+}$*MYC*$^{-}$ cells and the nearest *ICAM1*$^{+}$*CD68*$^{+}$ macrophage in three different CLL LNs. Statistical comparisons were performed within each CLL case using two-sided Wilcoxon rank sum tests. **C)** Quantification of *LGALS9* expression in concentric shells surrounding clustered *MS4A1*$^{+}$*MYC*$^{+}$ CLL PC cells. Note that there is significantly higher *LGALS9* signal proximal (10-200 μm) to clustered *MS4A1*$^{+}$*MYC*$^{+}$ CLL PC cells compared to distal regions (200-500 μm). Values were normalized per clustered *MS4A1*$^{+}$*MYC*$^{+}$ cell to the inner shell (inner = 1.0) and summarized as mean ± SEM. Statistical comparisons between inner and outer shells were performed using two-sided paired Wilcoxon signed rank tests. **D)** Immunofluorescence confocal images of CLL LNs showing staining for GAL9, MYC, CD20, CD44, CD8, and DNA within and around PCs (dashed outline). Individual fluorescence channels are shown on the right. The inset merged image corresponds to a higher magnification view of the proliferation center shown in Figure 6L. Scale bars, 100 μm (left), 25μm (right, insets).

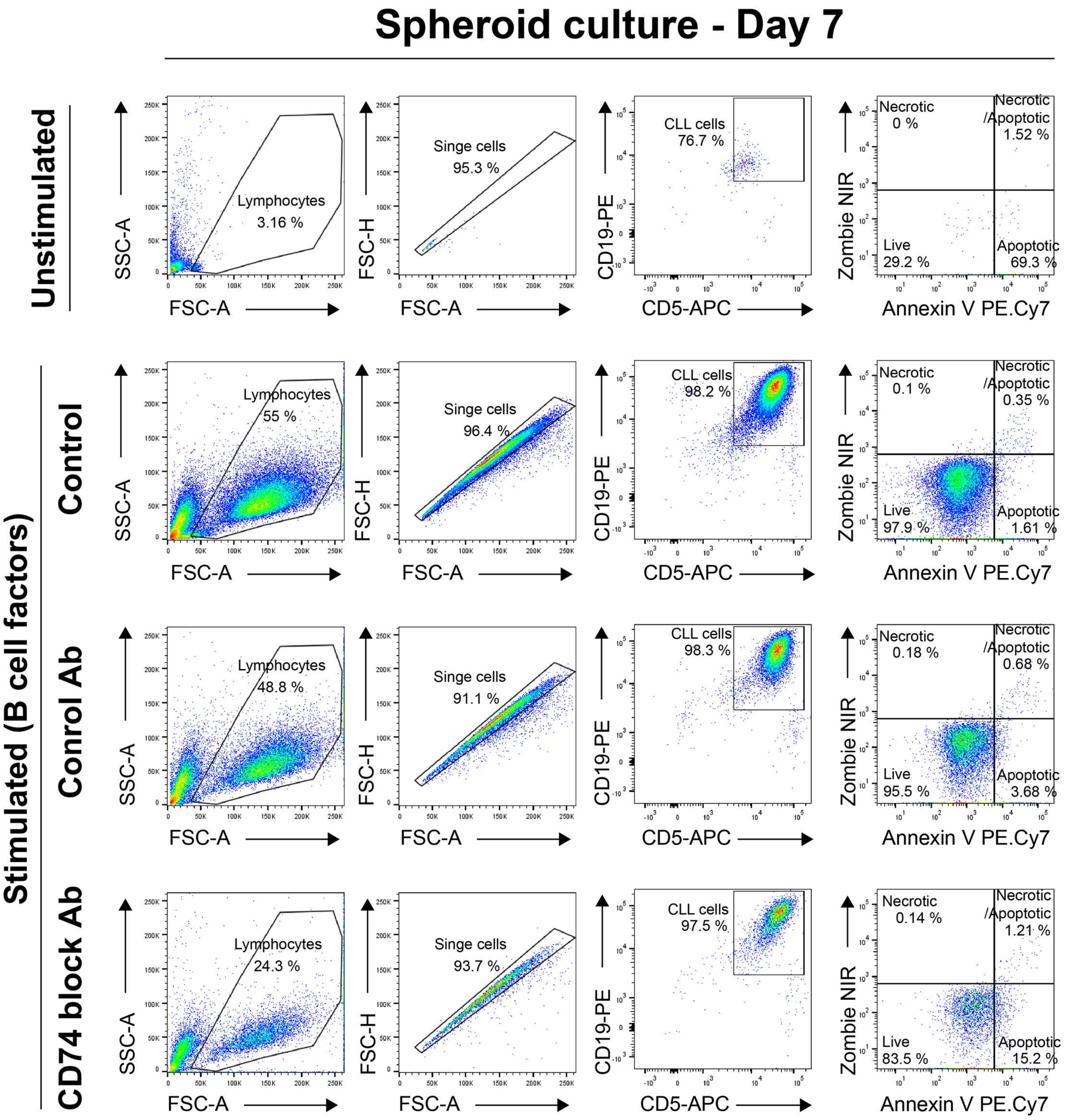


**Supplementary Figure 14. Flow cytometry gating of CLL cells in spheroid cultures.**

CLL cells isolated from peripheral blood were cultured as 3D spheroids for 7 days under the indicated conditions and analyzed by flow cytometry. A standardized gating strategy was applied across all samples: (i) lymphocyte gating based on FSC/SSC, (ii) singlet discrimination using FSC-A vs FSC-H, and (iii) CLL cell identification as $CD19^{+}CD5^{+}$ cells. Cell viability was assessed using Zombie NIR and Annexin V PE.Cy7 staining to delineate live, early apoptotic, necrotic, and late apoptotic/necrotic populations. The live $CD19^{+}CD5^{+}$ population gated here was used for CellTrace Violet proliferation analysis (shown in Figure 7E-F).
